# L-Fucose is a candidate monosaccharide neuromodulator and mitigates Alzheimer’s synaptic deficits

**DOI:** 10.1101/2022.08.11.503673

**Authors:** Jacopo Di Lucente, Jennyfer Tena, Ulises R. Mendiola, Xi Chen, Carlito B. Lebrilla, Izumi Maezawa, Lee-Way Jin

## Abstract

We identified a novel signaling function of L-fucose, a structurally unique monosaccharide. We showed that L-fucose enhanced excitatory neurotransmission and long-term potentiation (LTP) in Schaffer-collateral-CA1 synapses. L-fucose was released by neurons in an activity- and store-dependent manner, and induced rapid signaling changes to enhance presynaptic release. Such effects required L-fucose metabolism through the salvage pathway driven by fucokinase (FUK). Thus, L-fucose could be the first described monosaccharide neuromodulator affecting a metabolic-signaling mechanism. Human Alzheimer’s disease (AD) and 5xFAD mouse brains showed signs of fucose hypometabolism with impaired L-fucose signaling. Such abnormalities could be corrected by exogenous L-fucose, exemplified by rectification of LTPdeficits in 5xFAD hippocampus. Dietary L-fucose supplement, which increased cerebral free L-fucose levels and upregulated FUK to drive the salvage pathway, mitigated synaptic and behavioral deficits of 5xFAD mice. Our data reveals a therapeutic potential of oral L-fucose for AD, which is safe and easy to comply with.

## Introduction

Among monosaccharides, L-fucose is unique in its L-configuration whereas all other naturally occurring sugars in mammals have the D-configuration, and in its distinct structure lacking the hydroxyl group on the C-6 carbon. Its significance in mammalian physiology has been exclusively attributed to fucosylation – its incorporation into the oligosaccharide chains of *N*- and *O*-linked glycoproteins or glycolipid that confers functional modifications of macromolecules – a “fucosylation-centric” view^1^. In the brain, fucosylation was reported to affect processes involved in learning and memory, such as long-term potentiation, neurite outgrowth and migration, synapse formation, among others^1^. Fucosylation of synapsin I was found to regulate the turnover and stability of this key synaptic vesicle membrane protein regulating neurotransmitter release^2^. However, the exact mechanisms of how L-fucose, especially free L-fucose, affects brain function remain largely unknown.

Fucosylation in biological systems requires the generation of 5’-diphosphate-L-fucose (GDP-Fuc), a glycosyltransferase donor, through two pathways. The *de novo* pathway converts glucose-derived GDP-mannose to GDP-Fucose, and the salvage pathway utilizes free L-fucose to generate GDP-Fuc through an intermediate, fucose-1-phosphate (F-1P) via the rate-limiting enzyme fucokinase (FUK)^1^ (**Supplementary Fig. 1**). Greater than 90% of cellular GDP-Fuc is derived from the *de novo* pathway; the salvage pathway only plays a minor if not negligible role in replenishing cellular macromolecular fucose^3–5^. The function of the salvage pathway, particularly in the brain, is poorly understood although it would allow free L-fucose to influence cell functions. Indeed, our recent mass spectrometry study showed that differentiated neural cells efficiently incorporated ^13^C-labeled fucose intact into glycosylated macromolecules, suggesting an active salvage pathway in the brain^6^. In addition, hippocampal neurons released L-fucose in an activity-dependent manner (**Fig. 4a**), implying that free L-fucose may be involved in neural cell function via cellular or intercellular signaling.

The current study tests the hypothesis that free L-fucose modulates synaptic plasticity via activating the salvage pathway. We further asked whether such a mechanism is impaired in Alzheimer’s disease (AD), an archetypal disease of synaptic plasticity imposing huge impact on human society. Our findings suggest that L-fucose is a novel signaling molecule able to regulate synaptic neurotransmission via a metabolic signaling mechanism, and, contrary to the fucosylation-centric view, not likely via changing fucosylation of macromolecules. Furthermore, our data suggest that a fucose hypometabolism state links the triggers of AD to downstream synaptic deficits, which could be remedied by L-fucose supplementation to drive fucose metabolism via the salvage pathway.

## Results

### L-Fucose improves short-term presynaptic plasticity and long-term potentiation in 5xFAD hippocampus

We perfused hippocampal slices obtained from 12 months-old wild-type (WT) and 5xFAD mice with L-fucose (200 µM) and recorded the field excitatory postsynaptic potential (fEPSP) in Schaffer-collateral-CA1 synapses. In WT slices, the fEPSP increases were apparent within 1-2 min after initiation of perfusion, reached a plateau at ∼10 min, and returned to baseline ∼5 min after cessation of L-fucose (**Fig. 1a**). As controls for the specificity of L-fucose, three naturally occurring structurally close monosaccharides, D-fucose, the enantiomer of L-fucose, D-galactose that differs from D-fucose by having a hydroxyl group at C-6, and D-mannose that differs from D-galactose by inverting the stereochemistry of the hydroxyl groups at C-2 and C-4, did not alter fEPSP, suggesting L-fucose being a specific substrate for synaptic facilitation (**Fig. 1a and supplementary Fig. 2**). By comparison, fEPSP facilitation in 5xFAD slices was not elicited, even after prolonged perfusion. However, increasing the concentrations of L-fucose appeared to be able to switch on the responses of 5xFAD slices, albeit with substantial delays and subdued rates of increase (**Fig. 1b and supplementary Fig. 2**).

**Figure 1.**
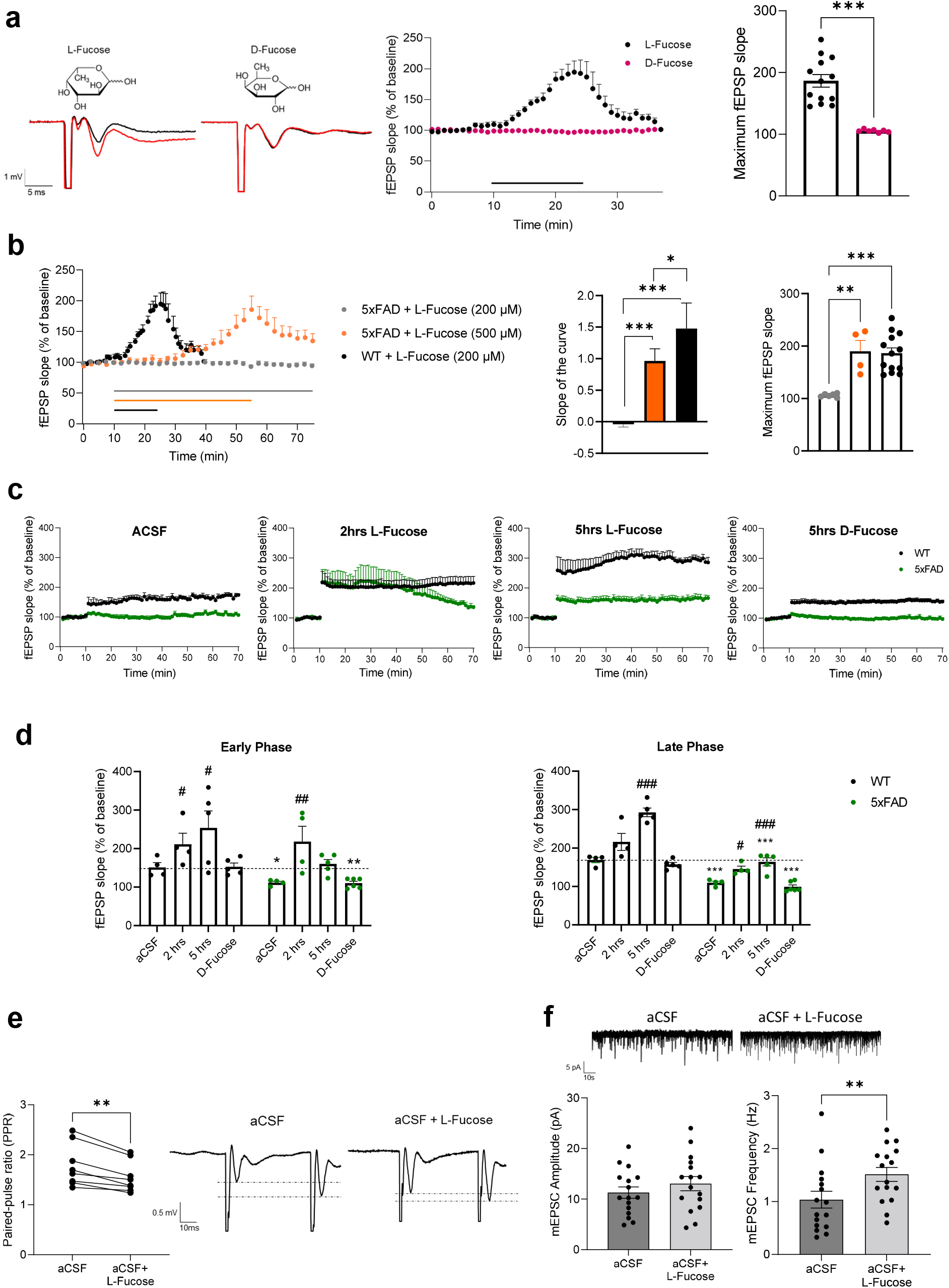
L-fucose modulates excitatory neurotransmission. All data are presented as means ± s.e.m.. **(a)** fEPSP recording in the stratum radiatum of hippocampal slices from WT mice perfused with 200 μM of L-fucose or D-fucose in aCSF. Representative traces before (black) and after (red) perfusion is shown. Scatter plot and quantification of maximum fEPSP slope values show that fEPSP slope is enhanced by L-fucose (*n*=14), but not D-fucose (*n*=7). *** *p*<0.001, Student’s unpaired t-test. **(b)** Stunted responses of 5xFAD hippocampus to L-fucose. Same data from WT in **1a** is plotted here for comparison with 5xFAD+L-fucose (200 μM) (*n*=6) and 5xFAD+L-fucose (500 μM) (*n*=6). \**p*<0.05, \*\**p*<0.01, \*\*\**p*<0.001; one-way ANOVA with Tukey’s *post hoc* test. **(c-d)** Pre-incubation with L-Fucose for 2 or 5 hours enhances hLTP in WT and 5xFAD hippocampi. Shown are scatter plots of hLTP (**c**) and quantification of fEPSP slope during the early and late phases of hLTP (**d**). *N*=4-5 per group. \**p*<0.05, \*\**p*<0.01, \*\*\**p*<0.001 compared to WT; #*p*<0.05, ##*p*<0.01, ###*p*<0.001 compared to respective aCSF control; one-way ANOVA with Tukey’s *post hoc* test. (**e**) L-fucose induces a decrease in paired-pulse ratio (PPR). *N*=8; ** *p*<0.01; Student’s paired t-test. (**f**) L-fucose increases the frequency, but not the amplitude, of mEPSCs in CA1 pyramidal neurons. Shown are representative tracings and the average amplitude and frequency values of mEPSCs over a 5 min period starting at 10 min of L-Fucose perfusion. *N*=16, ** *p*<0.05, Student’s paired t-test.

We further tested the effects of L-fucose on hippocampal long-term potentiation (hLTP), a cellular correlate of learning and memory, deficits of which are considered a precursor or substrate of dementia^7^. We and others previously showed hLTP deficits in 5xFAD mice in Schaffer-collateral-CA1 synapses following high frequency stimulation^8^. L-fucose, upon 2 hr pre-incubation, rectified hLTP deficits in 5xFAD hippocampus to approximating the WT level, although appearing not able to sustain the later phase (**Fig. 2c and 2d**). Interestingly, upon 5 hr preincubation, L-fucose further enhanced hLTP in WT hippocampus, but did not further enhance hLTP in 5xFAD hippocampus. This effect of L-fucose is specific as D-fucose showed no hLTP-enhancing effect (**Fig. 2c and 2d**). Taken together, both short-term facilitation and LTP studies suggest a deficit in the L-fucose metabolic pathway in 5xFAD brains. The ability of higher concentrations of L-fucose to partially rectify such a deficit suggests that L-fucose or its metabolites could drive forward its metabolic pathway.

**Figure 2.**
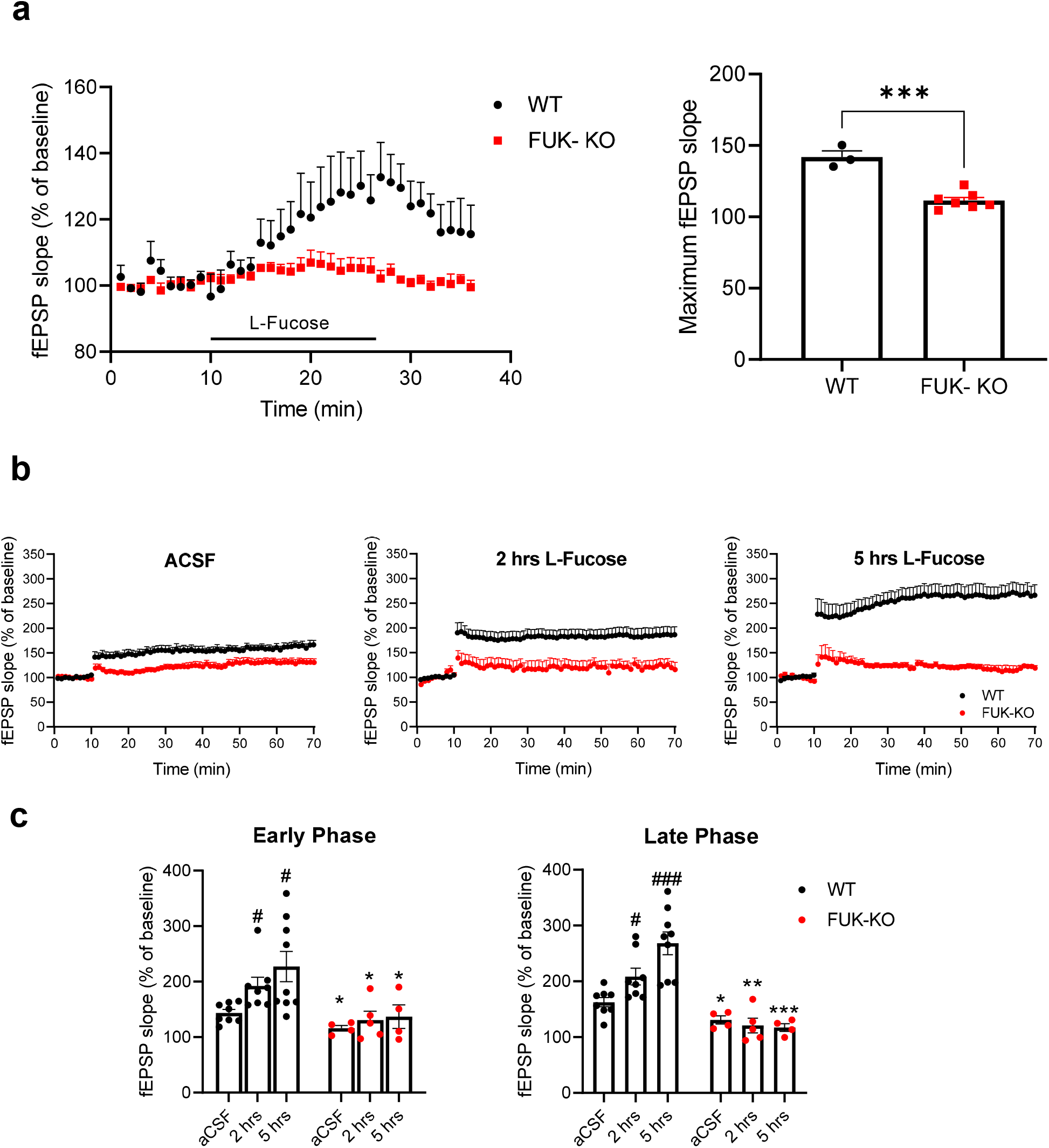
L-fucose effects depend on FUK. All data are presented as means ± s.e.m.. **(a)** Scatter plot and quantification of maximum fEPSP slope values show that fEPSP slope is enhanced by L-fucose (200 μM) in WT (*n*=3), but not in FUK-KO hippocampus (*n*=5). *** *p*<0.001, Student’s unpaired t-test. **(b-c)** Pre-incubation with L-Fucose does not enhance hLTP in FUK-KO hippocampi. Shown are scatter plots of hLTP (**b**) and quantification of fEPSP slope during the early and late phases of hLTP (**c**). *N*=4-5 per group. \**p*<0.05, \*\**p*<0.01, \*\*\**p*<0.001 compared to WT; #*p*<0.05, ###*p*<0.001 compared to respective aCSF control; one-way ANOVA with Tukey’s *post hoc* test.

To elucidate how L-fucose may facilitate neurotransmission, we conducted paired-pulse facilitation, a form of short-term synaptic plasticity, in WT hippocampal slices. The L-fucose-enhanced evoked EPSP (eEPSP) amplitude was associated with a decrease in the paired-pulse ratio (PPR) (**Fig. 1e**). Furthermore, L-fucose perfusion increased the frequency, but not the amplitude, of the miniature excitatory postsynaptic current (mEPSC) (**Fig. 1f**). These results indicate that the L-fucose effects were mediated by changes in release probability of presynaptic glutamatergic synaptic vesicles, and not by postsynaptic mechanisms. To evaluate whether the above defective synaptic responses to L-fucose in 5xFAD mice can be generalized to other AD mouse strains, we conducted the short-term facilitation experiment using 12 month-old APP-PS1 mice^9^. Similar to 5xFAD mice, hippocampal slices from APP-PS1 mice showed no response to 200 μM L-fucose but a subdued and delayed response to 500 μM L-fucose (**Supplementary Fig. 3**).

### L-Fucose enhances synaptic plasticity via driving the salvage pathway

Regarding the metabolic mechanisms via which L-fucose enhances synaptic plasticity, the salvage pathway is particularly relevant as it utilizes free L-fucose derived from dietary sources, added to culture medium/perfusate (as in our experiments), or salvaged from degradation of glycoproteins (**Supplementary Fig. 1**)^10, 11^. We further ruled out the involvement of the *de novo* pathway as 6-alkynyl-fucose (6-Alk-Fuc), a specific and potent inhibitor of GDP-4-keto-6-deoxymannose 3,5-epimerase (FX)^12^ that catalyzes the last step of the *de novo* pathway, did not affect L-fucose-facilitated fEPSP (**Supplementary Fig. 4**). We therefore hypothesized that L-fucose affects synaptic plasticity via driving the salvage pathway. To evaluate this hypothesis, we tested the effects of L-fucose in Fcsk ^em1(IMPC)Bay^ mice deficient in fucokinase (FUK), the gatekeeping enzyme that converts L-fucose to F-1P in the salvage pathway^1^ (**Supplementary Fig. 1**). FUK knockout (FUK-KO) abolished the short-term facilitation effect of L-fucose (**Fig. 2a**) and significantly attenuated hLTP (**Fig. 2b and c**). In contrast to WT or 5xFAD mice, the application of L-fucose did not augment hLTP, either by 2 hr or 5 hr pre-incubation (**Fig. 2b and c**). Taken together, our data suggest that the synaptic effects of L-fucose are mediated by the salvage pathway, requiring the function of FUK.

### Reductions of L-fucose and defective salvage pathway in AD brain tissue

We next tested the hypothesis that the salvage pathway is defective in AD in view of the subdued responses to L-fucose in 5xFAD hippocampus. qPCR showed that transcripts of FUK and FPGT, the two enzymes constituting the salvage pathway (**Supplementary Fig. 1**), were reduced in 5xFAD (**Fig. 3a**) and pathologically confirmed human AD brains (**Fig. 3b and Supplementary Table 1**) compared to age-matched controls. In contrast, GMD and FX, the two enzymes constituting the *de novo* pathways, were not significantly different between AD and control samples, except that FX was found increased in human AD brains. Western blotting of FUK further confirmed reduced FUK protein levels in both 5xFAD (**Fig. 3c**) and human AD samples (**Fig. 3d**). Furthermore, the levels of free L-fucose were significantly reduced in AD samples (**Fig. 3e**). Notably, in 5xFAD hippocampal slices, the FUK protein level was enhanced in a time-dependent manner by L-fucose treatment, reaching the WT level following 5 hr treatment, suggesting a positive regulation of FUK by L-fucose (**Fig. 3f**). This set of data indicates both a deficiency of available free L-fucose and a defective salvage pathway to metabolize free L-fucose in AD brains. The rescue effects of applied L-fucose, albeit being subdued, could be due to its ability to supplement the deficient free L-fucose in 5xFAD brains and drive forward the salvage pathway in a limited capacity, in part via increasing the expression of FUK.

**Figure 3.**
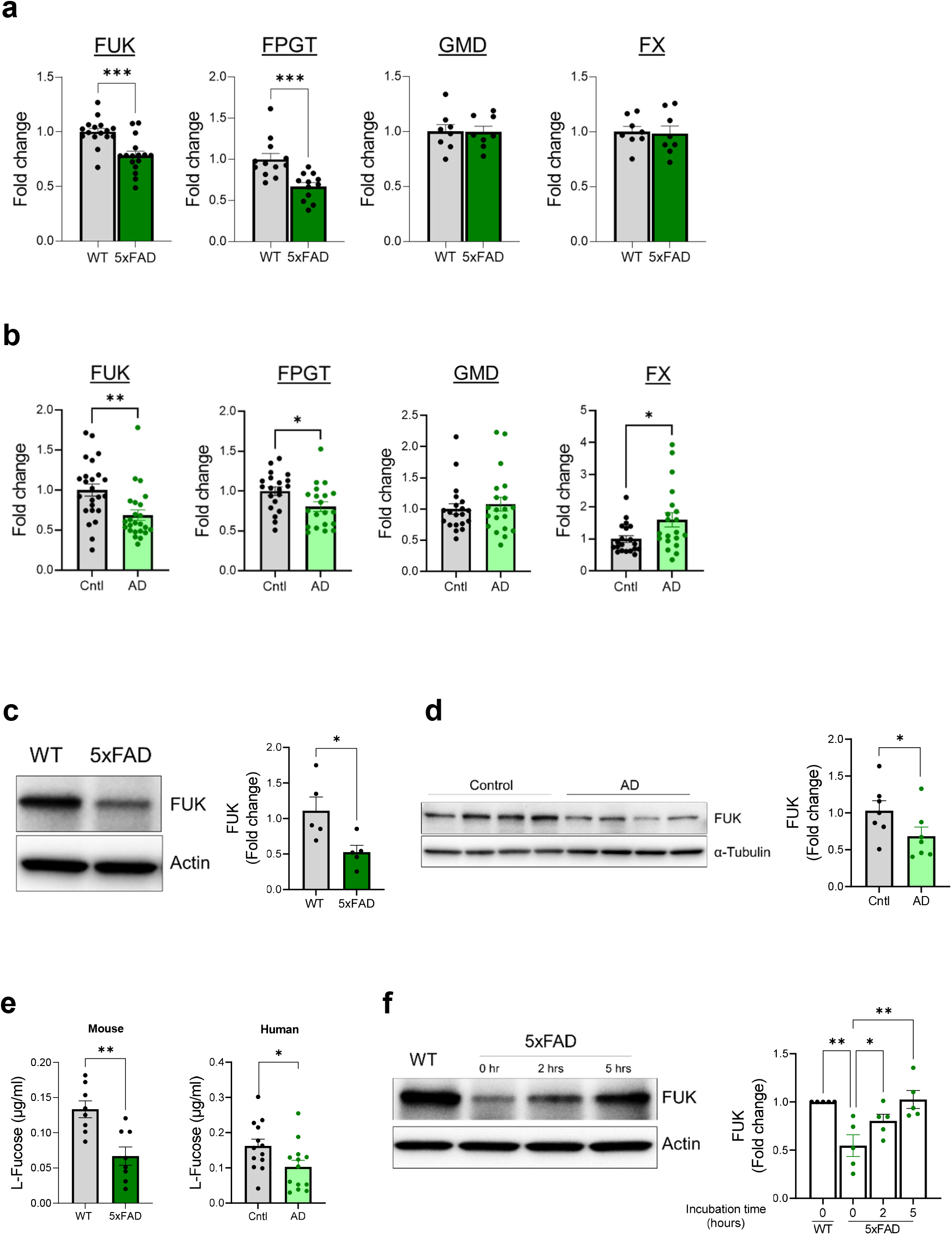
Abnormal L-fucose metabolism in 5xFAD and human AD brain tissue. All data are presented as means ± s.e.m.. **(a-b)** Quantitative PCR shows that transcript levels of the enzymes in the salvage pathway but not in the *de novo* pathway are decreased in **(a)** 5xFAD mice (*n*=16/group, \*\*\**p*<0.001 Student’s unpaired t-test), and in **(b)** human AD brains (*n*=20/group, *p<0.05, **p<0.01 Student’s unpaired t-test). **(c-d)** Western blotting shows that protein levels of FUK are decreased in **(c)** 5xFAD mice (*n*=5, \**p*<0.05; Student’s unpaired t-test) and in **(d)** middle temporal cortex of human AD brains (*n*=6, *p<0.05 Student’s unpaired t-test). **(e)** Free L-fucose levels are decreased in hippocampal slices of 5xFAD mice (*n*=8/group) and middle temporal cortex of human AD brains (*n*=13/group). \**p*<0.05, \*\**p*<0.01, Student’s unpaired t-test. **(f)** Western blotting shows that L-fucose time-dependently increases FUK levels in 5xFAD hippocampi. Shown are a representative Western blot and quantitative data compiled from 5 independent experiments; \**p*<0.05, \*\**p*<0.01, one-way ANOVA with Tukey’s *post hoc* test.

### The synaptic effects of L-fucose are not likely fucosylation-dependent

The dependence of the L-fucose effects on the salvage pathway may support the assumption that L-fucose affects synaptic transmission via fucosylation of glycoconjugates, as GDP-Fuc, the end product of the salvage pathway, is the substrate utilized by all fucosyltransferases. However, this is not compatible with the time frame of its actions as new glycosylation is a co- and post-translational event that takes hours^13^. As shown above, the short-term effects of L-fucose occurred within 1-2 minutes of application (**Fig. 1**), prompting us to question the involvement of fucosylation. Moreover, preincubation with 6-alkynyl-fucose, a potent inhibitor of cellular fucosylation^12^, did not block the short-term facilitation of L-fucose (**Supplementary Fig. 4**). To validate this notion, we evaluated changes of fucosylation by lectin blots. Following up to 5 hours of perfusion of WT or 5xFAD hippocampal slices with L-fucose, we did not find any significant changes of fucosylation detected by lectins Ulex Europaeus Agglutinin 1 (UEA-1), Aleuria Aurantia Lectin (AAL), and Len Culinaris Agglutinin (LCA) that are specific for either terminal or core fucosylation (**Supplementary Fig. 5**). As lectins may not detect minor fucosylation changes, we further perfused hippocampal slices with ^13^C-fucose to facilitate detection of newly incorporated fucose moiety in an N-glycome analysis using sensitive LC-MS/MS methods^14^. Up to 5-hour perfusion, we could not detect incorporation of ^13^C-fucose into any N-glycans. Moreover, in the comprehensive representation of all fucosylated-only (without sialic acid moiety) N-glycans and sialofucosylated N-glycans, no significant difference of fucosylation was observed within 5 hours of treatment, nor between WT and 5xFAD slices (**Supplementary Fig. 6**). These results strongly suggest that the synaptic effects of L-fucose are not dependent on changes in fucosylation, but more likely rely on much quicker signaling events.

### L-fucose is a candidate endogenous signal molecule/neuromodulator

The observation that free L-fucose is present in both human and mouse brains in moderate abundance (**Fig. 3e**) and facilitates fEPSP within seconds of application suggests that it may function as a yet unrecognized neuromodulator and, together with its intracellular metabolites, serve signaling functions within or between cells. To further test this possibility, we asked the following questions pertaining to key properties of a neuromodulator: 1) whether L-fucose is released by neurons in an activity-dependent manner; 2) whether the release can be regulated by intracellular stores of L-fucose; and 3) whether L-fucose can induce immediate and early signaling changes in neural cells in a short time frame that is clearly not contingent upon fucosylation. Using differentiated primary hippocampal neurons, we found that following KCl-induced depolarization, L-fucose release was increased in a time-dependent manner, culminating at 10 min before subsiding toward the baseline (**Fig. 4a**), suggesting that endogenous L-fucose can be released following activity. We next hypothesized that L-fucose-containing glycoconjugates serves as a cellular store from which L-fucose could be released. To modulate cellular L-fucose levels, we preincubated the differentiated primary hippocampal neurons in cultures with deoxyfuconojirimycin (DFJ), a potent inhibitor of α-L-fucosidase known to enrich cellular fucose store^15^. Incubation with DFJ prior to KCl-induced depolarization accelerated the release of L-fucose, which culminated at 1 min, and further sustained it at high levels for at least one hour. DFJ treatment without KCl did not result in changes of L-fucose release (**Fig. 4a**). We next determined if such an enhanced release of endogenous L-fucose would facilitate neurotransmission in hippocampal slices similar to externally applied L-fucose. Preincubation with DFJ indeed induced short-term facilitation in WT hippocampus comparable to external application of L-fucose. As a negative control, the DFJ effect was not observed in 5xFAD slices, consistent with deficient 5xFAD responses to endogenous L-fucose (**Fig. 4b**). Interestingly, this result implies that cellular L-fucose present in glycoconjugate forms could be released extracellularly or into the synapse by an unknown mechanism that does not involve fucosidases.

**Figure 4.**
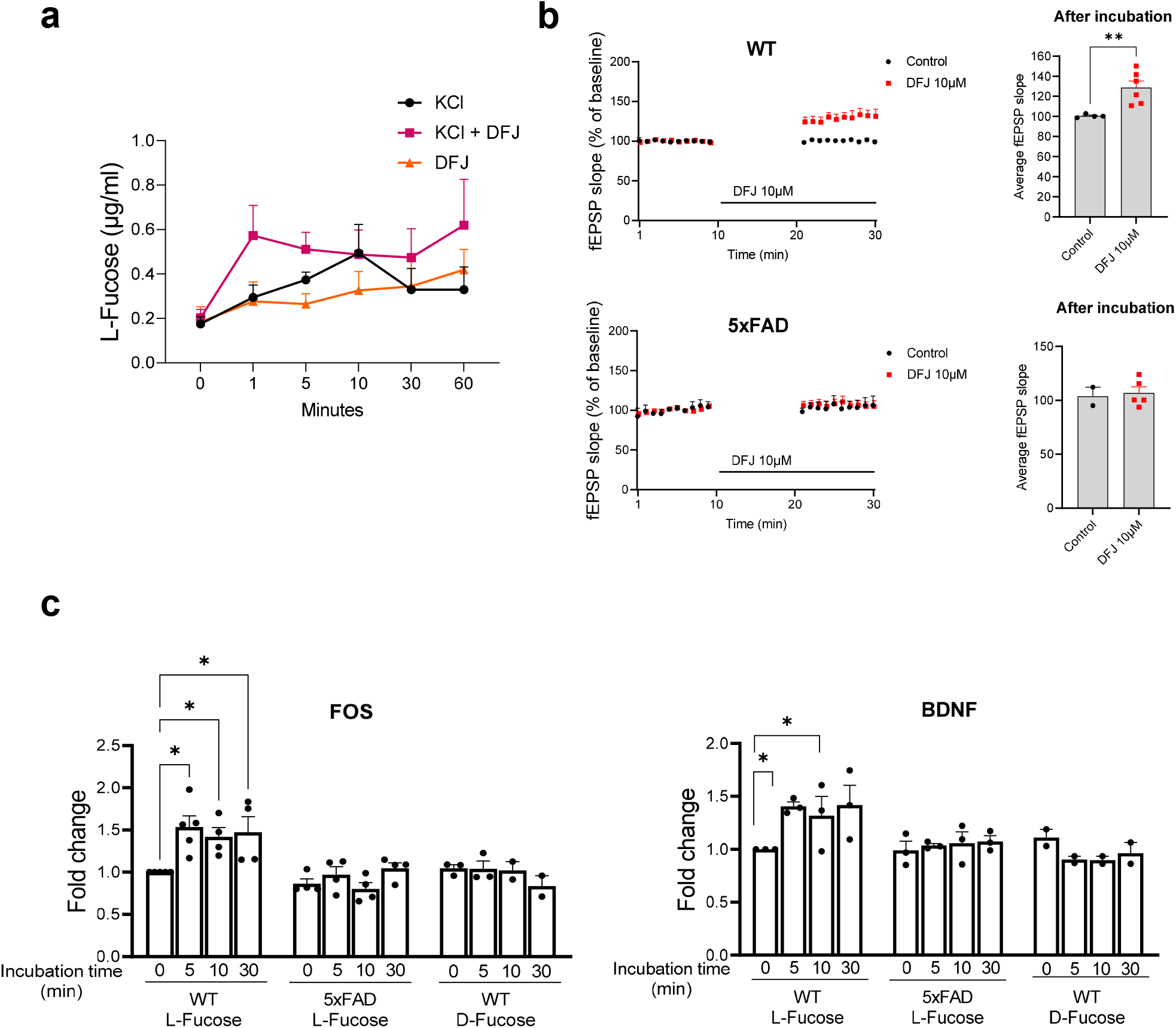
L-fucose shows activity- and store-dependent release and elicits rapid signaling changes. All data are presented as means ± s.e.m.. **(a)** Neuronal depolarization by KCl (300 mM) induces L-fucose release, which is further enhanced by pre-incubation with DFJ (10 µM) for 10 minutes before KCl. Shown are L-fucose levels in the conditioned media of cultured well-differentiated mouse hippocampal neurons. *N*= 4-5/group. Two-way ANOVA with Tukey’s *post hoc* test shows a significant difference in released L-fucose levels by type of stimulation (F(2)=5.165, *p*=0.009) and by time (F(5)=2.788,*p*=0.025). A post-hoc test revealed significant pairwise differences between KCL+DFJ and KCl alone (*p*=0.013) and between KCL+DFJ and DFJ alone (*p*=0.012). There are also significant differences among time points. Compared to 0 min, the values at 1 min (*p*=0.019), 10 min (*p*=0.025), 30 min (*p*=0.047) and 60 min (*p*=0.020) show significant differences. Comparisons between values at 1 min show a significant difference between KCL+DFJ and KCL alone (*p*=0.046). **(b)** DFJ enhances the slope of fEPSP recorded in the stratum radiatum of WT hippocampal slices (*n*=4-6), but not that of 5xFAD slices (*n*=2-5). The scatter plot shows fEPSP slopes in a 10-min baseline recording which was discontinued during the 10-min preincubation with DFJ (10 μM), and resumed for the following 10 min. Bar graphs present average fEPSP slopes in the last 10 min of recording. \*\**p*<0.01, Student’s unpaired t-test. **(c)** L-fucose (but not D-fucose) increases the transcript levels of *fos* and *bdnf* within 5 min of treatment in WT hippocampal slices but not in 5xFAD slices. *N*=3-4/group; \**p*<0.05; one-way ANOVA with Tukey’s *post hoc* test.

We next conducted experiments over short time frames to measure several key activity-related mediators of neuronal plasticity. Treating WT hippocampal slices with L-fucose rapidly (within 5 min) increased the RNA levels of the *c-fos* and *bdnf*, two neuronal activity-induced immediate early genes implicated in synaptic plasticity^16, 17^ (**Fig. 4c**). This transcriptional effect is specific as *c-Jun* or *Jun-B* was not activated (**Supplementary Fig. 7**). In contrast, D-fucose treatment of WT slices or L-fucose treatment of 5xFAD slices failed to induce c-fos or *bdnf* expression (**Fig. 4c**). Because our data indicate that L-fucose enhances presynaptic release, we next examined activation of Ca^2+^/calmodulin-dependent protein kinase II (CaMKII) and phosphorylation of its substrate synapsin I, key events leading to presynaptic release^18^. L-fucose induced phosphorylation of both CaMII and synapsin I, which was apparent within 5 min of application (**Supplementary Fig. 8a-c**). Other signaling molecules that were quickly activated by L-fucose include CREB, but not ERK (**Supplementary Fig. 8d and e**). In contrast, 5xFAD hippocampus showed significantly subdued responses (**Fig. 4c and Supplementary Fig. 8**). Taken together, we showed that L-fucose is endogenously released in an activity- and store-dependent manner, and rapidly activates signaling pathways to modulate synaptic plasticity, making it a candidate neuromodulator.

### Dietary supplement of L-fucose mitigates AD-like synaptic deficits in 5xFAD mice

Our results suggest that administration of L-fucose could supplement brain L-fucose to drive the fucose salvage pathway in 5xFAD mice and mitigate AD-like defects in synaptic plasticity and cognition. To test this possibility, we allowed 11 month-old 5xFAD and WT littermate mice to consume 0.5% L-fucose-supplemented diet (FD) or control diet (CD, same diet composition without L-fucose supplement) ad lib for 30 days. Mice receiving FD were well groomed and displayed no observable differences, including body weight and food consumption habit, compared to those receiving CD (**Supplementary Fig. 9**). L-fucose supplement significantly increased the level of free L-fucose in the 5xFAD brains to a level comparable to WT (**Fig. 5a**). It also improved their exploratory behavior, evidenced by increased traveling distance in the open field (**Fig. 5b**), and mitigated their memory deficits, evidenced by improved performance in novel object recognition (**Fig. 5c**) and T-maze tests (**Fig. 5d**). Furthermore, we tested the abnormal sensorimotor gating mechanisms noted in prodromal stages of AD^19^ using acoustic response tests. These tests examine the response to sudden, loud noises, as measured by acoustic startle response (ASR) and prepulse inhibition (PPI). We found that CD-fed 5xFAD mice had reduced magnitude and prolonged latency of ASR to loud stimuli (120 dB) compared to CD- or FD-fed WT mice (**Fig. 5e**). While WT animals showed inhibition of the startle response with soft but increasingly loud prepulse tones, the inhibition was significantly reduced in CD-fed 5xFAD mice (**Fig. 5f**). This PPI impairment suggests inability to filter out unnecessary information due to abnormal sensorimotor gating mainly from entorhinal lesions^19, 20^, previously demonstrated in humans with early stage of AD^21^. FD consumption significantly alleviated the ASR and PPI deficits of 5xFAD mice (**Fig 5e and f**).

**Figure 5.**
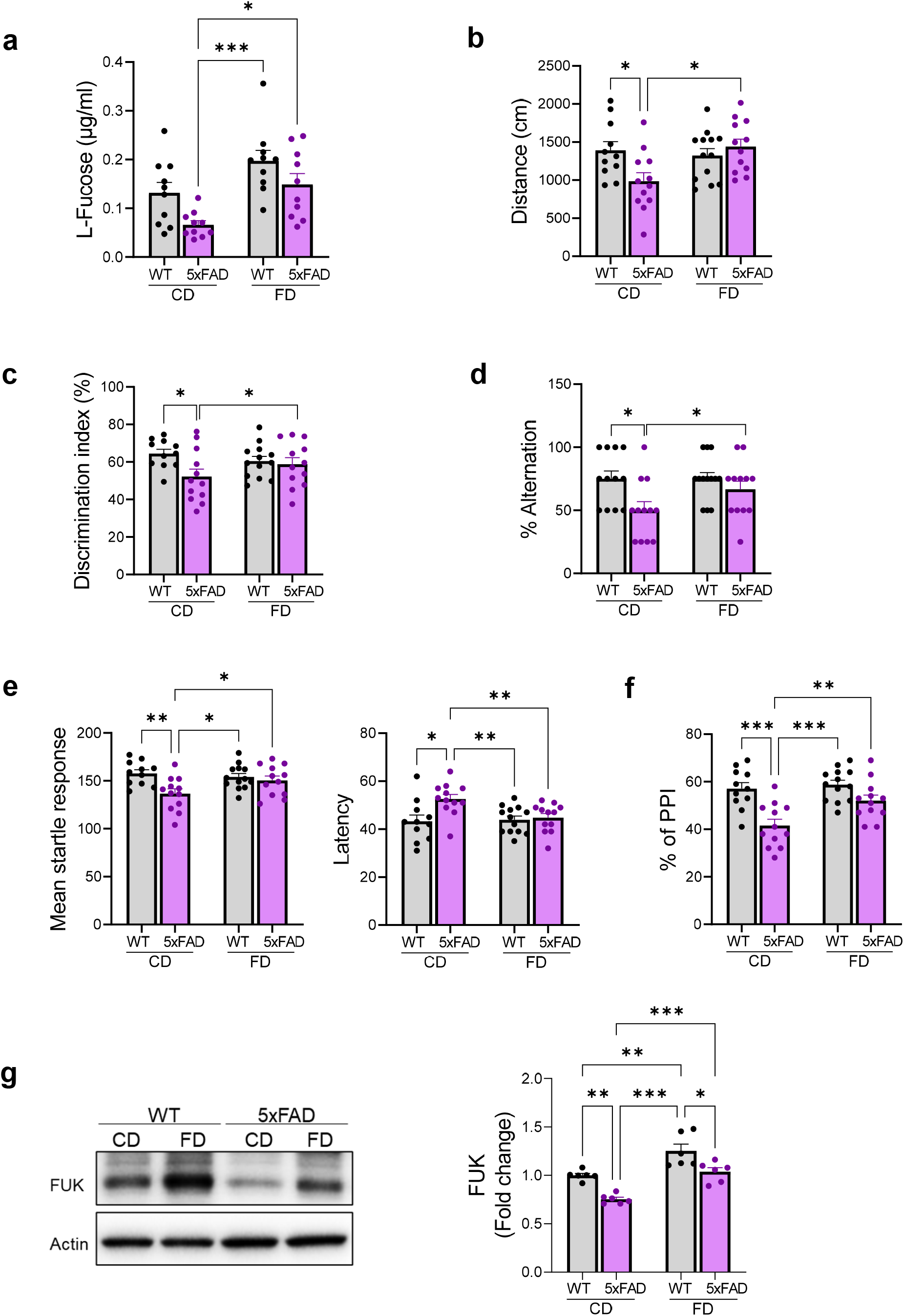
L-fucose supplement mitigates behavioral and synaptic deficits of 5xFAD mice. Eleven months-old WT and 5xFAD mice were assigned to FD or CD for 30 days. All data are presented as means ± s.e.m.. **(a)** FD increases the L-fucose levels in 5xFAD forebrain to the WT levels. *N*=10/group. **(b-f)** FD improves exploratory, memory, and sensorimotor gating deficits of 5xFAD mice: **(b)** distance traveled in cm in open field test; **(c)** percentage of discrimination index in novel object recognition test; **(d)** percentage of alternation in T-maze test; **(e)** mean startle response and latency (in ms) in response to a 120 dB startle stimulus; and **(f)** percentage of prepulse inhibition using a 70 dB prepulse followed by a 120 dB startle stimulus. *N*=11 for WT-CD, 12 for 5xFAD-CD, 13 for WT-FD, and 12 for 5xFAD-FD. **(g)** Western blotting shows that FD increases the FUK protein levels in 5xFAD forebrain to the levels of WT and further increases its levels in WT mice. *N*=6/group. \**p*<0.05, \*\**p*<0.01, \*\*\**p*<0.001, two-way ANOVA with Tukey’s *post hoc* test.

Corresponding to the memory improvement, dietary supplementation of L-fucose also enhanced hLTP in 5xFAD mice, particularly in the late phase (**Supplementary Fig. 10**). This effect of L-fucose in hLTP appeared to require FUK. First, FUK-KO mice fed CD showed substantially reduced magnitude of hLTP compared to WT mice in both induction and maintenance. Second, such stunted hLTP was not improved by FD consumption, indicating that FD-dependent hLTP enhancement was mediated by the FUK-catalyzed salvage pathway (**Supplementary Fig. 11**). In congruence with *in vitro* data, FD consumption by 5xFAD mice also resulted in increased levels of FUK (**Fig. 5g**), supporting the notion that administered L-fucose could supplement the deficient free L-fucose in 5xFAD brains and drive the salvage pathway to mitigate AD-like synaptic deficits. We further showed that FD consumption also rectified several key molecules in synaptic plasticity, such as activation of synapsin I, CaMKII, and CREB (**Supplementary Fig. 12**). Such effects occurred in the absence of detectable upregulation of fucosylation, analyzed by lectin blots (**Supplementary Fig 13a**). Flow cytometry analyses using fucosylation-specific lectins as well as an antibody to Lewis X antigens (CD15, a series of carbohydrate epitopes with terminal fucosylation and abundantly expressed in neural tissue^22, 23^) further supported this assessment (**Supplementary Fig. 13b-d**), indicating that dietary L-fucose supplementation did not significantly increase fucosylation of brain macromolecules. Furthermore, the effects of dietary L-fucose supplementation were not related to a reduction of AD-like amyloid pathology as it did not reduce the levels of cerebral amyloid deposition or Aβ peptides (of both soluble and insoluble forms) in 5xFAD mice (**Supplementary Fig. 14**), supporting a direct action of L-fucose on synaptic plasticity.

## Discussion

We identified a novel signaling function of L-fucose that is not likely dependent on fucosylation of macromolecules, which makes an exception to the dominant fucosylation-centric view of L-fucose function^1, 4^. We further showed that this function depends on its metabolism through FUK in the salvage pathway, suggesting a metabolic signaling mechanism. Based on its rapid release by neurons in an activity- and store-dependent manner and its rapid induction of signaling changes enabling presynaptic glutamate release, L-fucose is qualified as a candidate neuromodulator, the first described in the chemical category of sugars. Based on our data, we propose a model illustrated in **Fig. 6**. This model expands the complex mechanisms regulating excitatory neurotransmission. L-fucose may have dual impacts on synapses: short-term signaling independent of fucosylation as described here, and long-term modifications of glycoproteins via fucosylation that affect turnover of synaptic proteins, neurite outgrowth, and neuronal architecture, as described by Hsieh-Wilson and colleagues^2^. Both modes of action appear to converge on synapsin I; its phosphorylation can be quickly induced by free L-fucose, and its fucosylation in the form of fucose-α(1-2)-galactose protects it from degradation and thereby preserves neuronal architecture^2^. Due to our limited understanding, our model does not address the possible non-cell autonomous functions of L-fucose to influence neighboring neurons, astrocytes, and microglia. Possible broader signaling effects of free L-fucose on other types of synapses or other physiological processes are also not well understood.

**Figure 6.**
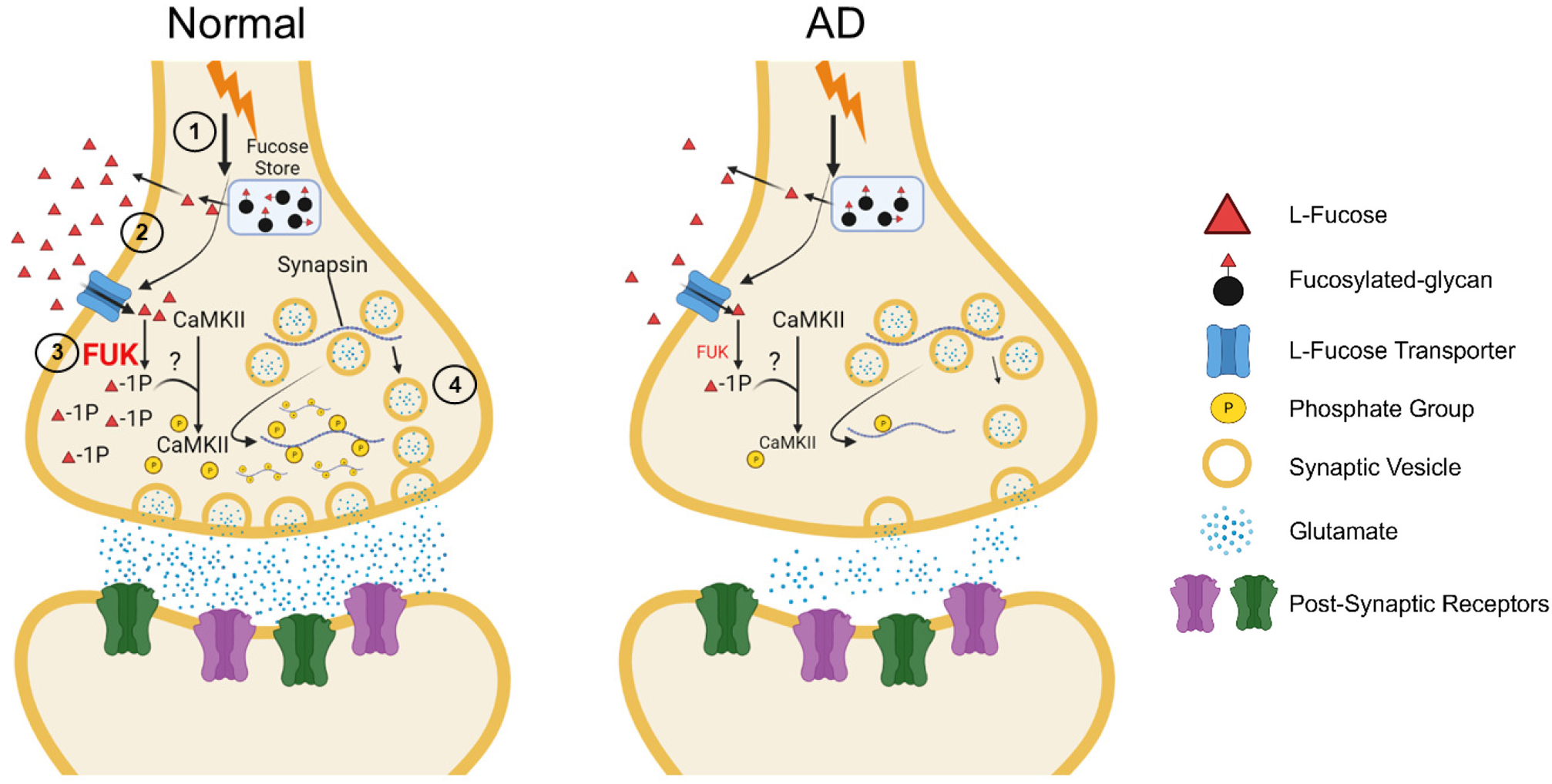
A working model of L-fucose actions in normal and AD synapses. The actions are summarized in four steps: **(1)** Activity- and store-dependent L-fucose release. Neuronal activity, such as depolarization or action potential, induces L-fucose release from an internal store composed of fucosylated glycans. **(2)** Internalization of L-fucose by fucose transporter. Cell-released or diet-derived L-fucose is internalized by a fucose transporter via facilitated diffusion to be the substrate for the salvage pathway. **(3)** FUK-dependent metabolic signaling. FUK is required for L-fucose effects and generates fucose-1-phosphate or other metabolites that activate signaling pathways involving CaMKII, c-fos, CREB, BDNF, and synapsin I. **(4)** Enhancement of presynaptic glutamate release. Phosphorylation of synapsin I contributes to the increased release probability of presynaptic glutamatergic synaptic vesicles, thus enhancing fEPSP and hLTP. In AD, free L-fucose level, FUK expression, and levels of phosphorylated CaMKII and synapsin I are decreased, resulting in compromised regulation of excitatory neurotransmission.

Prior reports from two groups implicated the signaling function of L-fucose. Pioneering work by Krug and colleagues showed that exogenously applied L-fucose enhanced hippocampal LTP *in vitro*^24, 25^, which was replicated by us using mouse models. They speculated that L-fucose interfered with the interactions between synaptic receptors or ion channels, many of which are glycoproteins, and endogenous lectins. Recently Wu and colleagues showed that exogenously applied L-fucose rapidly activated Wnt signaling in CHO cells^26^. However, the study only used free L-fucose as a competitive probe to support a putative receptor that would bind fucosylated LRP-6 (a Wnt co-receptor) and regulate Wnt complex endocytosis. The possible signaling function of free L-fucose was not stipulated. Notably, neither group described L-fucose effects as physiological, and both attributed the effects to “competitive interference” of glycoprotein function in a manner conforming to the fucosylation-centric view.

Although the competitive interference model remains possible, it is not likely based on our observation that the effects of L-fucose depend on its intracellular salvage pathway metabolism. Furthermore, L-fucose is rapidly released by neurons in an activity- and store-dependent manner and rapidly induces signaling pathways to affect synaptic plasticity, suggesting it being an endogenous physiological neuromodulator. Our findings call for further studies of its mechanisms of action. One may consider that L-fucose, like many neuromodulators, acts as a ligand recognized by cellular receptors or sensors. It is known that L-fucose can serve as an energy substrate for the microbiota, which have developed well-regulated L-fucose sensing systems that control their growth and virulence^27^. So far there is no evidence supporting the existence of such L-fucose receptors or sensors in mammalian cells. Alignment studies for fucose permease in *Campylobactor jejuni*^28^ and FusK and FusR in *Escherichia coli*^29^, all well-studied bacterial fucose sensors, did not yield homologous genes or proteins in mammalian cells (Dr. Chang-en Yu, personal communication).

On the other hand, a cell surface L-fucose transporter that internalizes L-fucose by facilitated diffusion was partially purified from the mouse brain^30, 31^. Considering the requirement of FUK for L-fucose effects, we propose a metabolic signaling mechanism of L-fucose (**Fig. 6**), similar to that of glucose. Glucose, following intracellular transport, is rapidly phosphorylated by hexokinases into glucose-6-phosphate (G-6P) (**Supplementary Fig. 1**). Hexokinases are rate-liming enzymes of glucose metabolism while its downstream metabolites including G-6P are signaling molecules^32, 33^. In a parallel fashion, dietary or cell-released L-fucose is rapidly internalized and phosphorylated by the rate-liming enzyme FUK into F-1P, which we speculate is a signaling metabolite. We hypothesize that FUK, similar to hexokinases, serves as a “signaling nexus” that integrates metabolic and activity cues to regulate downstream signaling^34^. The ability of L-fucose (or its metabolites) to quickly upregulate FUK is supportive of this notion.

Emerging evidence supports that synaptic hypometabolism, manifested by imaging evidence of reduced glucose uptake, precedes synaptic dysfunction and imposes a risk of future cognitive decline and AD^35^. While most attention has been focused on glucose, our data show signs suggesting fucose hypometabolism in AD. We found that free L-fucose level and its salvage pathway enzymes were significantly reduced in human AD and mouse 5xFAD brains, a condition that limits efficient usage of L-fucose. We further found that increasing cellular fucose store by fucosidase inhibition resulted in enhanced activity-induced L-fucose release and more robust fEPSP (**Fig. 4**), suggesting that releasable and functional L-fucose can be drawn from internal macromolecular fucose. As macromolecular fucose is largely derived from glucose via the *de novo* pathway (**Supplementary Fig. 1**), it is possible that abnormal glucose metabolism in AD may indirectly limit the availability of free L-fucose and the capacity of activity-dependent L-fucose release, particularly at the synapse, contributing to fucose hypometabolism. Our model suggests that fucose hypometabolism disrupts the cerebral “metabolic wiring” and contributes to synaptic dysfunction in AD (**Fig. 6**).

Exogenous L-fucose in free or conjugated forms may have multiple health benefits. In particular, fucoidans, which are polysaccharides rich in seaweeds and characterized by high L-fucose and sulfate contents, are suggested to have therapeutic potentials in multiple diseases, including AD^36, 37^. Our previous study found that in human milk oligosaccharides (HMO), L-fucose is the most abundant component, a major distinguishing feature between human and bovine milk^38^. The superior health benefits of HMO have been attributed in part to their ability to regulate the gut microbiota^39^. While the microbiota-gut-brain axis could provide an indirect mechanism of how L-fucose could affect brain function, a more direct mechanism would be that free L-fucose, in the diet or generated from catabolism of fucosylated glycans (in HMO or fucoidans) by intestinal microflora and subsequently up-taken by the colon^4^ and passing the blood-brain barrier^40^, could directly affect the brain. Our FD study lends support to this mechanism as FD consumption remedied the low levels of cerebral free L-fucose in 5xFAD mice, enhanced hLTP, and improved performance in tasks enabled by multiple functional synaptic systems. Furthermore, parallel to upregulation of FUK by L-fucose incubation of hippocampal slices, FD consumption enhanced the cerebral FUK level to drive the salvage pathway. Our data leads to a therapeutic hypothesis that oral consumption of L-fucose can improve synaptic function in AD via mitigating fucose hypometabolism. Because L-fucose consumption is safe and easy to comply with, it is a candidate therapy that can be readily tested in human clinical trials.

## Online Methods

### Mice

Tg6799 5xFAD [B6SJL-Tg(APPSxFILon,PSEN1*M146L*L286V)6799Vas/Mmjax] mice on the C57Bl/6 background were originally obtained from Dr. Robert Vassar, Ph.D., Northwestern University^41^. APP-PS1 [B6 .Cg-Tg(APPswe,PSEN1 d E9) 85Dbo/Mmjax] mice on the C57Bl/6 background were originally purchased from the Jackson Laboratory. WT littermates were used as WT controls. Fcsk ^em1(IMPC)Bay^ mice were obtained from Arthur Beaudet, Ph.D., Baylor College of Medicine, and generated for the Knockout Mouse Production and Phenotyping Project (KOMP2). The strain, C57BL/6N-*Fcsk^em^*^1^*^(IMPC)Bay^*/Mmmh, is available from the MMRRC, RRID:MMRRC_068158-MU. Phenotypic details can be found on the International Mouse Phenotyping Consortium site at https://www.mousephenotype.org/data/genes/MGI:1916071. Mice were randomly assigned to treatment groups and phenotyping parameters, including neurobehavioral assessments were evaluated in a blinded manner. Roughly equal numbers of male and female mice were used. All protocols involving mouse models were approved by the Institutional Animal Care and Use Committee.

### Human brain samples

All tissues were collected and provided by the University of California Davis Alzheimer’s Disease Research Center (P30 AG072972). Written informed consent, including consent for autopsy, was obtained from study participants or, for those with substantial cognitive impairment, a caregiver, legal guardian or other proxy. Study protocols were reviewed and approved by the Institutional Review Board (IRB). For postmortem diagnosis, we followed the National Institute on Aging-Alzheimer’s Association guideline for the neuropathologic assessment of AD^42^. The brain tissues used for the current study were snap frozen during autopsy and was stored in −80°C before use. Patient characteristics are summarized in Supplementary Table 1.

### Chemicals and designed diets

L-Fucose, D-Fucose, D-Galactose, and D-Mannose were purchased from Sigma-Aldrich (Sigma, St. Louis, MO). 6-Alkynyl-L-Fucose was purchased from Fisher Scientific (Thermo Fisher Scientific, Waltham, MA). For diet treatment, control diet (TD. 140368) and 0.5% L-Fucose supplemental diet (TD. 200724, which has the same composition as the control diet with the addition of 5 g/kg L-fucose and 0.1 g/kg red food color for identification) were designed by a certified Teklad nutritionist at Envigo (Indianapolis, IN).

### Brain slice preparation

Brain slices containing the hippocampus were obtained as previously described^43^. The mice were anesthetized with isoflurane and decapitated. The brain was rapidly removed and submerged in ice-cold high-sucrose artificial cerebrospinal fluid containing (in mM): 220 sucrose, 2 KCl, 0.2 CaCl_2_, 6 MgSO_4_, 26 NaHCO_3_, 1.3 NaH_2_PO_4_, and 10 d-glucose (pH 7.4, set by aeration with 95% O_2_/5%CO_2_). Coronal brain sections (300 μm thick) containing the dorsal hippocampus were cut in ice-cold modified ACSF using a DTK-1000 D.S.K Microslicer (Ted Pella, Inc., Redding, CA, USA). Slices were then immediately transferred to a nylon net submerged in normal ACSF containing (in mM): 126 NaCl, 3 KCl, 2 CaCl_2_, 1 MgCl_2_, 26 NaHCO_3_, 1.25 NaH_2_PO_4_, and 10 d-glucose (pH 7.4, set by aeration with 95% O_2_/5% CO_2_), for at least 40 min at 35 °C controlled temperature. After subsequent incubation for at least 1 h at room temperature, hemi-slices were transferred to a recording chamber, submerged in normal ACSF with a constant flow of ∼ 2 ml/min.

### Extracellular recording of fEPSPs, pair pulse facilitation, and long-term potentiation

Field EPSPs were evoked after stimulation of the Schaffer collateral afferents by a bipolar stimulating electrode (WPI, Sarasota, FL), and recorded from the stratum radiatum of the CA1 region using a borosilicate capillaries filled with 3M NaCl (resistance 1-3 MΩ). Baseline stimulation rate was 0.05Hz. fEPSPs were filtered at 2 kHz and digitized at 10 kHz with a Multiclamp 700B amplifier (Molecular Devices, Sunnyvale, CA). Data were collected and analyzed with pClamp 10.3 software (Molecular Devices). Slope values of fEPSPs were obtained for quantitation of the responses. Paired-pulse facilitation (PPF) of synaptic transmission was induced by electrical stimulation of Schaffer collaterals with paired pulses at an inter-stimulus interval of 50 ms. The PPF was quantified as the ratio of the second fEPSP slope over that of the first response. For LTP, after 10 minutes of stable baseline recording of fEPSPs evoked every 20s at a constant current pulse of 0.2-0.4 mA with a duration of 60 µs, high-frequency stimulation (HFS), consisting of two trains of 100-Hz (1s) stimulation with the same intensity, was applied. Recording was then continued for 60 min with stimulation of fEPSPs every 20 s.

### Whole-cell patch-clamp recordings

Whole-cell patch-clamp recording was conducted as previously described^44^. To record miniature excitatory post-synaptic currents (mEPSCs), patch pipettes (5-8 mΩ) were filled with 270 mOsm internal solution containing the following (in mM): 120 K-gluconate,10 KOH, 9 KCl, 3.48 MgCl2, 4 NaCl, and 10 mm HEPES. AMPA receptor currents were recorded from cells voltage clamped at −70 mV, in bath solution containing 10 μM bicuculline (to block GABAA receptors) and 1 μM TTX (to block presynaptic action potentials). Bath application of 1,2,3,4-tetrahydro-6-nitro-2,3-dioxo-benzo[f]quinoxaline-7-sulfonamide disodium salt (NBQX, 10 μM) was used as a control to confirm that currents were attributable to AMPA receptor activity. Analysis of mEPSCs was performed manually using Mini analysis software (Synaptosoft, Inc., version 6.0.2) with a noise amplitude threshold of 2 pA.

### L-fucose quantification

Measurement of L-Fucose levels was conducted using the L-Fucose Assay kit from Megazyme (Megazyme, Wicklow, Ireland). This kit quantifies the level of L-fucose via this reaction catalyzed by L-fucose dehydrogenase (L-FDH): L-Fucose + NADP^+^--------> L-fucono-1,5-lactone + NADPH + H^+^. For brain homogenates, samples were deproteinized by using the Deproteinizing Samples Preparation kit (BioVision, Milpitas, CA). For conditioned media from hippocampal neuronal cultures, aliquots were incubated at 95⁰C for 10 minutes to inactivate enzyme activity. After centrifugation, supernatant was use for the assay. Ten µl samples were added to a 96-well plate with distilled water, solution I (buffer), and solution II (NADP+), provided by the manufacturer. A full calibration curve was used for this assay. Absorbances of the solution was read at 340 nm after approximately 4 min and the reaction was started by the addition of 5 µl of L-FDH solution. Absorbance was read again for 10 minutes in kinetic mode.

### Tissue homogenate preparation and Western blotting

Brain tissues were homogenized in lysis buffer (150 mM NaCl, 50 mM Tris-HCl, 2 mM EDTA, 1% Triton X-100, 0.5% SDS, 0.1% Na-deoxycholate) with protease inhibitor cocktail and phosphatase inhibitor (Sigma). Equivalent amounts of protein were analyzed by 4%–20% Tris-Glycine gel electrophoresis (Invitrogen). Proteins were transferred to polyvinylidene difluoride (PVDF) membranes and probed with antibodies. Visualization was enabled using enhanced chemiluminescence (GE Healthcare Pharmacia). The following primary antibodies (dilutions) were used: anti-FUK (1:1000, Invitrogen), anti-phospho-Synapsin I (1:1000, Cell Signaling), anti-Synapsin I (1:1000, Cell Signaling), anti-phospho-CaMKII (1:1000, Cell Signaling), anti-CaMKII (1:1000, Cell Signaling), anti-phospho-CREB (1:1000, Cell Signaling), ant-CREB (1:1000, Cell Signaling) and Actin (1:5000, Cell Signaling). Secondary antibodies were HRP-conjugated anti-rabbit or anti-mouse antibody (1:1000, Cell Signaling).

### Quantitative real-time PCR

Total RNA from tissue samples were extracted using RNeasy® Plus Universal Mini Kit (Qiagen, Valencia, CA) according to manufacturer’s protocol. cDNA was synthesized using iScript Reverse Transcription Supermix (Bio-Rad, Hercules, CA). RNA purity and concentrations were assessed by measuring the absorbance at 260 nm, and 280 nm through a NanoDrop 2000C Spectrophotometer (Thermo Scientific, Waltham, MA). Quantitative PCR (qPCR) was performed using the Sso Fast EvaGreen Supermix (Bio-Rad) in the CFX96 Touch Real-Time PCR Detection System (Bio-Rad). Mouse and human primers were purchased from Bio-Rad. Gene expression was normalized to an endogenous reference gene, β-actin. Data were analyzed by the 2^-ΔΔCt^ method. All experiments were performed in duplicate.

### Primary hippocampal neuron culture

Primary hippocampal neuron cultures derived from newborn C57BL/6J mice were prepared as previously described^45^. Cells were plated on poly-d-lysine-coated 6 wells plate at a density of 5 × 10^5^ cells per well in serum-free Neurobasal medium plus B27 supplement and maintained at 37°C in 5% CO_2_.

### N-glycome analysis

**(1) Cell membrane extraction from hippocampus tissue.** Hippocampal samples were homogenized and resuspended in homogenization buffer containing 0.25 M sucrose, 20 mM HEPES-KOH (pH 7.4), and a 1:100 protease inhibitor cocktail. Cells were lysed on ice using a probe sonicator operated with alternating on and off pulses of 5 and 10 s, respectively. Lysates were pelleted by centrifugation at 9000xg for 10 min to remove the nuclear fraction and cell debris. The supernatant was transferred to high-speed tubes, loaded onto a Beckman Optima TLX Ultra-centrifuge at 4°C, and centrifuged at 200,000 x g for 45 min in series to remove other nonmembrane subcellular fractions. The resulting cell membrane pellet was stored at - 20°C until further processing. Optimizations to this approach has been shown to generate a purified cell membrane pellet^14^. **(2) Enzymatic release and purification of N-glycans.** Proteins were suspended with 100 µL of 100 mM NH_4_HCO_3_ in 5 mM dithiothreitol and heated at 100°C for 10s to thermally denature the proteins. To release the glycans, 2 µL of peptide N-glycosidase F were added to the samples, followed by incubation in a microwave reaction at 60°C for 10 min to accelerate N-glycans release. Samples were incubated for 18 h at 37°C to hydrolyze the N-glycans. The reaction was quenched with 350 μL of water followed by ultracentrifugation at 200,000 x g to separate the N-glycans and the membrane fraction (MF)containing our lipids and de-glycosylated proteins. The released N-glycans were purified by solid-phase extraction using porous graphitized carbon (PGC) packed cartridges. The cartridges were first equilibrated with nanopure water and a solution of 80% (v/v) acetonitrile and 0.05% (v/v) trifluoroacetic acid in water. The dried samples were solubilized, loaded onto the cartridge, and washed with nanopure water to remove salts and buffer. N-Glycans were eluted with a solution of 40% (v/v) acetonitrile and 0.05% (v/v) trifluoroacetic acid in water, dried and reconstituted in 30µl of water prior to mass spectrometric analysis. **(3) Glycomics analysis by LC-MS/MS.** Purified mouse brain N-glycans were analyzed using an Agilent nano-LC/chip Q-ToF MS system. The nano-LC system employs a binary solvent consisting of A (0.1% formic acid in 3% acetonitrile in water (v/v)) and B (0.1% for-mic acid in 90% acetonitrile in water (v/v)). Samples were enriched and separated on the Agilent HPLC-Chip comprised of a 40 nL enrichment column and a 75 μm x 43 mm ID analytical column both packed with porous graphitized carbon in 5 μm particle size. The sample was delivered by the capillary pump to the enrichment column at a flow rate of 3 μL min^-1^ and separated on the analytical column by the nano-pump at a flow rate of 0.3 μL min^-1^ with a gradient that was previously optimized for N-glycans: 0% B, 0-2.5 min; 0-16% B, 2.5-20 min; 16-44% B, 20-30 min; 44-100% B, 30-35 min; and 100% B, 35-45 min followed by pure A for 20 min of equilibration. MS spectra were acquired at 1.5 s per spectrum over a mass range of m/z 600–2000 in positive ionization mode. Mass inaccuracies were corrected with reference mass m/z of 1221.991. N-Glycan compositions were identified using MS and MS/MS data as well as an in-house retrosynthetic library based on the mammalian N-glycan biosynthetic pathway. Deconvoluted masses were compared to theoretical masses using a mass tolerance of 20 ppm and a false discovery rate of 0.5% on the Agilent MassHunter software version B.7. Relative abundances were determined by integrating peak areas for observed glycan masses and normalizing to the summed peak areas of all glycans detected.

### Open-field test

The SmartCage^TM^ system (29.8 x 18 x 12.8 cm) (AfaSci, Inc., Redwood City, CA) was used to assess spontaneous activity in homecages. Automated data analysis was performed using CageScore^TM^ software (AfaSci, Inc.). The home cage activity was determined by infrared beam breaks (x, y, and z photo-beam break counts). Mice were allowed to explore freely for 10 min. Movements of mice were recorded by Cage Center software and analyzed by Cage Score 2 software as described^46, 47^.

### Novel object recognition test

Novel object recognition protocol was similar to previously described^43^. Briefly, the apparatus consisted of a white plastic arena (40 x 40 cm) placed in the experimental room under low lighting conditions (∼20 lux). The mouse was habituated to the arena without objects for 10 minutes for 2 consecutive days. On the third day, after habituation, the mouse was placed back to the home cage and the arena was cleaned with ethanol. Two identical objects, which were predetermined to have no natural significance or association with the control mice, were secured to the bottom of the arena. The mouse was allowed to explore the objects for 10 minutes, and then placed in the home cage again. The arena and objects were clean with ethanol. One same object and one different object were placed into the arena. The mouse was returned to the arena for a 10-minute exploration of the same and novel objects, which was recorded and the discrimination index percentage was calculated.

### T-Maze spontaneous alternation test

Each mouse was subjected to 5 trials. The T-maze test apparatus had 3 arms (30 × 10 cm2) and was elevated 30 cm above the ground. One trial consisted of a choice and a sample run. During the choice run one of the two target arms was blocked by a barrier according to a pseudorandom sequence, with equal numbers of left and right turns per session and with no more than two consecutive turns in the same direction. The mouse was allowed to explore the accessible arm. Before the sample run (intertrial interval of ∼10 s), the barrier was removed enabling accessibility to both arms. On the sample run the mouse was replaced back into the start arm facing the experimenter. The mouse was allowed to choose one of the two target arms. The trial was classified as success if the animal chose the previously blocked arm.

### Acoustic startle reflex and PPI

An automated Startle Reflex System (SR-LAB; San Diego Instruments, Inc., San Diego, CA, USA) was used to measure acoustic startle reflex and PPI. Before the experiments, detailed piezoelectric sensitivity tests and dB calibration were performed to ensure reproducible startle responses. The system consisted of a startle chamber housed in a sound attenuated isolation cabinet equipped with an internal fan and light. A cylindrical transparent acrylic holding apparatus resting within the isolation chamber was used to hold each subject throughout the testing session. Background noise and acoustic stimuli were controlled through the SR-LAB microcomputer and interface assembly and kept constant at 65 dB. Mice were placed in cylinders 10 min prior to the initial startle stimuli and only background noise was offered during this acclimation period. To measure acoustic startle response and PPI, the five trials were: no stimulus, two types of startle stimulus only (100 and 120 dB, 20 ms broad band burst), and two types of startle stimulus preceded by a prepulse (a 20 ms broad band burst). The onset of the prepulse was separated from the startle onset by a 100 ms prepulse-startle interval (PSI), and the prepulse intensity used was 70 dB. Each was repeated nine times in random order. The trials were separated by an average interval of 30 s. SR-LAB software was used to calculate latency and startle response. PPI values were calculated as a percentage score for each prepulse using the following formula:

PPI(%)=100-[(ASR for prepulse + pulse trial)/(ASR for pulse alone trial)]*100.

### Flow cytometry

To assess brain cell fucosylation after L-fucose supplemented diet, hippocampi were dissociated enzymatically using the Neural Tissue Dissociation Kit (Miltenyi Biotec). Cells suspension was resuspended in PBS. Two µg/ml of AAL-Fluorescein, LCA-Fluorescein (Vector Laboratories, Newark, CA) was added, or monoclonal Mouse anti-Mouse SSEA-1 (against Lewis X/CD15, clone MC-480, at 1:200 dilution, LSBio, Seattle, WA), and samples were incubated at 37⁰C for 1 hour in the dark. After incubation, the cells were pelleted at 300 x *g* for 5 min at 4⁰C and washed with PBS. The cells were then resuspended in 3 ml PBS. Cell staining was excited at 488nm and the emission was detected via the FL1 channel by a flow cytometer, BD Accuri C6 Plus (BD Sciences). Data were analyzed using FlowJo 10.8 (BD Sciences).

### Human Aβ42 ELISA

Brain tissue samples from mice treated with L-fucose or control diet were fractionated into Tris-buffered saline (TBS)-soluble, and SDS-soluble fractions to assess the solubility of amyloid-β species as previously described^48^. Briefly, for the TBS-soluble fraction, tissues were homogenized in Tris-buffered saline (TBS) with protease inhibitors, followed by centrifugation at 100,00 x g for 1 hour at 4⁰C. The pellets were then homogenized in SDS buffer containing 2% SDS in TBS with protease inhibitors, incubated on ice for 10 min, followed by centrifugation at 100,000 x g for 1 hour at 4⁰C. Aβ42 levels were measured using Human Aβ42 ELISA Kits (Thermo-Fisher Scientific), following the manufacturer’s instructions.

### Immunofluorescence staining of amyloid

Freshly dissected brains were fixed in 10% formalin and 1-mm coronal slices containing hippocampus and cerebral cortex were embedded in paraffin. Six µm paraffin sections were obtained by microtome and used for immunofluorescence. Sections were dewaxed in Xylene and hydrated through alcohol gradient. For antigen recovery, slides were boiled in NaCitrate (10 mM, pH 6) for 15 min. After 1 hr incubation with 10% normal goat serum (Invitrogen) to block non-specific binding, sections were stained with the amyloid dye (E,E)-1-fluoro-2,5-bis(3-hydroxycarbonyl-4-hydroxy) styrylbenzene (FSB, Sigma) (500 nM) and counterstained with propidium iodide (500 nM). Images were taken using a SPOT RT sCMOS camera (SPOT Imaging, Sterling Heights, MI) and analyzed using the NIH ImageJ software.

### Statistical Analysis

Statistical analysis was performed using GraphPad Prism 9 software. All data are presented as means ± s.e.m. Comparison of the mean values was performed using unpaired Student’s two-tailed t-test, on-way ANOVA, or two-way ANOVA with Tukey’s *post hoc* test. Exact sample sizes and statistical test used for each comparison were provided in corresponding figure legends. *P* < 0.05 was considered to be statistically significant.

## Acknowledgements

We thank Dr. Chang-en Yu, University of Washington at Seattle U.S.A. for alignment studies in search of homologous genes or proteins of bacterial L-fucose receptors or sensors in mammalian cells; and Dr. Cheng-Chang Lien, Institute of Neuroscience, National Yang Ming Chiao Tung University, Taiwan, for advice. This work was funded by U.S. NIH grants R01 AG062240 (NIA) to I.M., L-W.J., and C.B.L.; and P30 AG072972 (NIA) to L-W.J..

## Author Contributions

I.M. and L-W.J. contributed equally and serve as co-corresponding authors. J.D. and I.M. performed major electrophysiological, cell culture, and biochemistry experiments; I.M. and L-W.J. conceived and designed the study; U.R.M. maintained the mouse colonies; C.B.L. designed the glycomics analysis and J.T. conduct it and analyze the data; X.C. provided quantitative and analytic supports; all investigators contributed to the planning of the experiments and writing of the manuscript.

## Supplementary Figure Legends

**Supplementary Figure 1.**
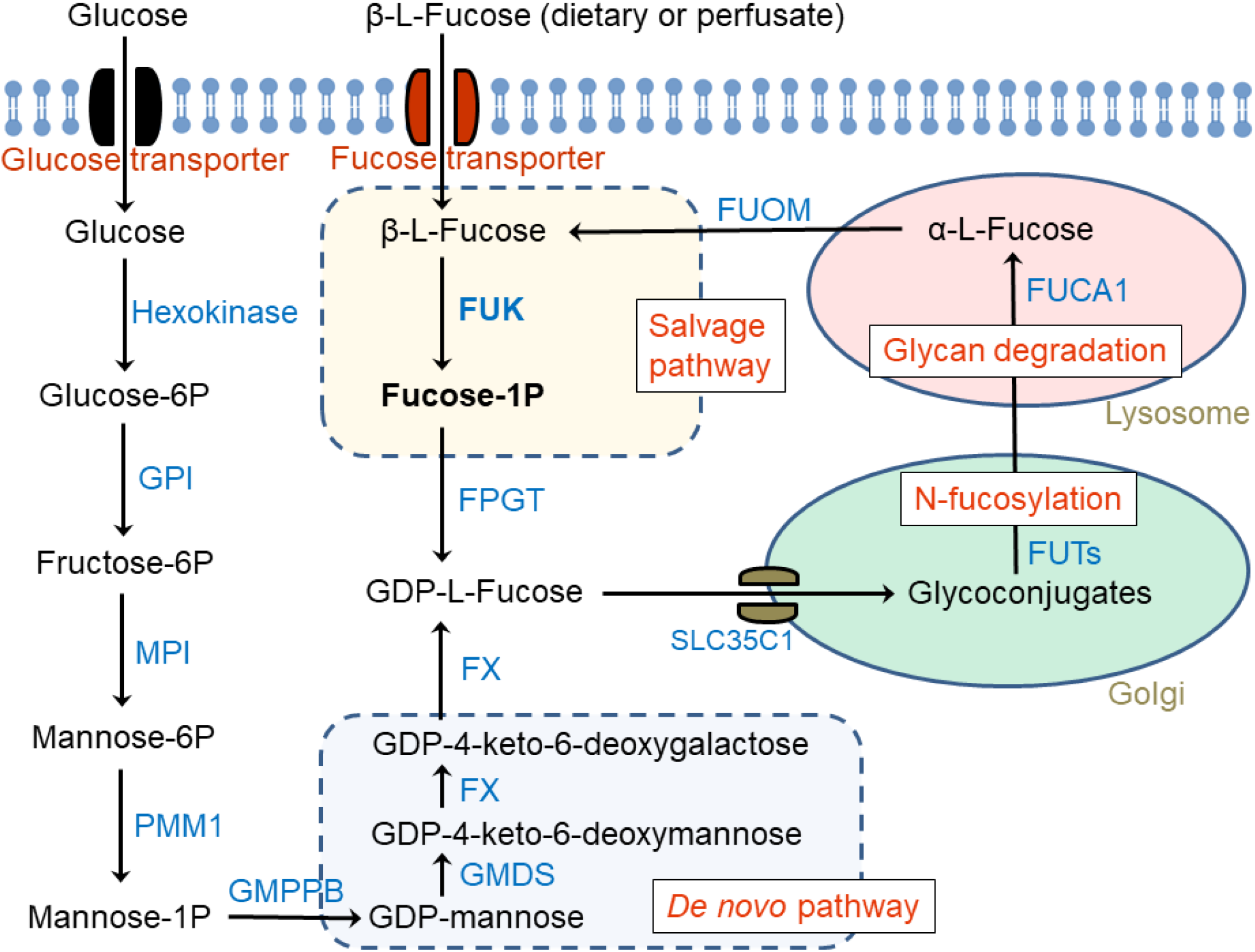
Fucose metabolism pathways. Shown are the two metabolic pathways for GDP-L-fucose synthesis by the salvage and *de novo* pathways. A part of glucose metabolism is highlighted to draw a parallel to the salvage pathway metabolizing internalized L-fucose and to illustrate its influence on GDP-L-fucose synthesis via the *de novo* pathway. FPGT, fucose-1-phosphate guanylyltransferase; FUCA1, α-L-fucosidase 1; FUK, fucokinase; FUOM, fucose mutarotase; FUTs, fucosyltransferases; FX, GDP-L-fucose synthase; GMDS, GDP-mannose 4,6-dehydratase; GMPPB, GDP-mannose pyrophosphorylase B; GPI, glucose-6-phosphate isomerase; MPI, mannose phosphate isomerase; PMM1, phosphomannomutase 1.

**Supplementary Fig. 2.**
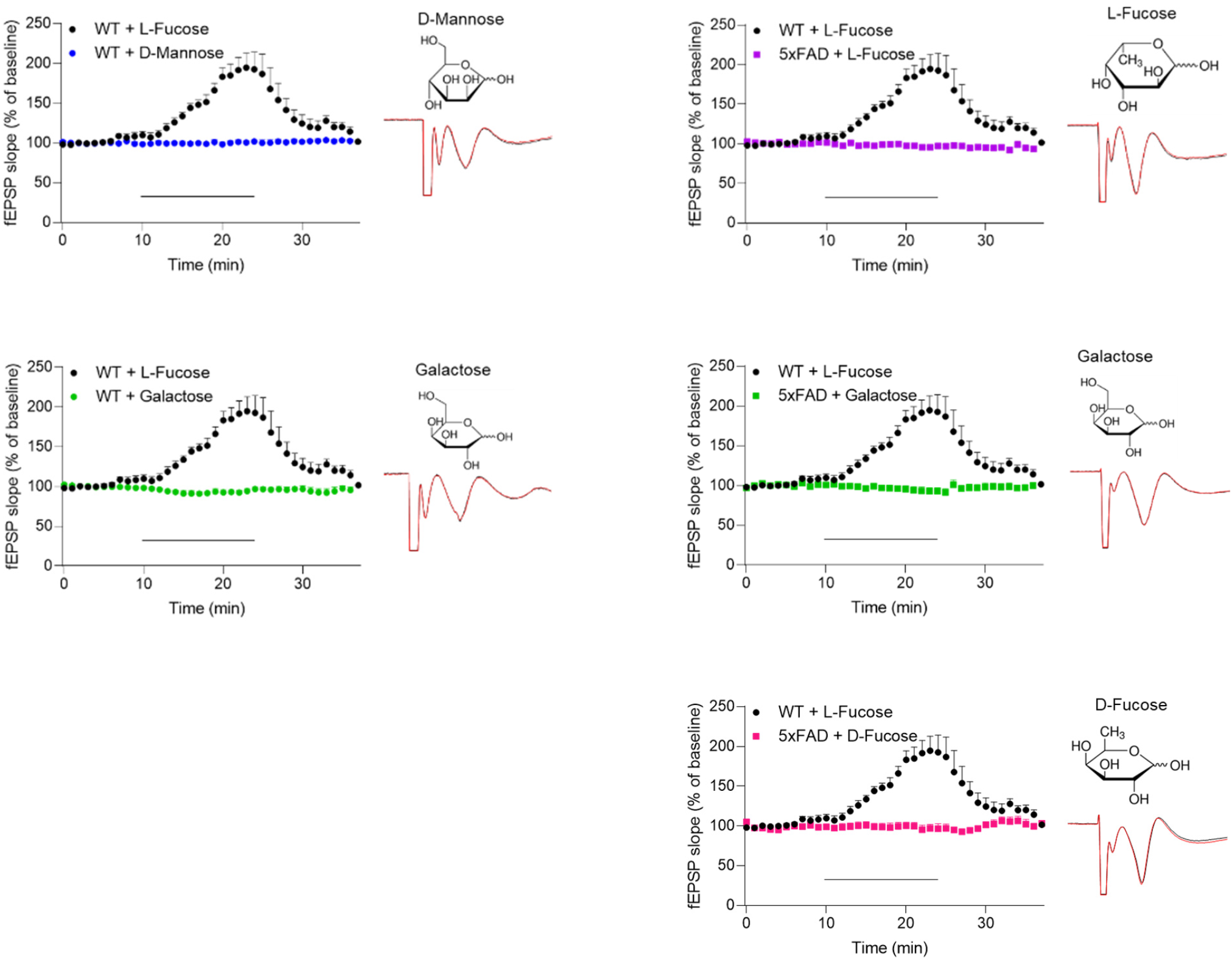
L-fucose effects are specific. fEPSPs of Schaffer-collateral-CA1 synapses were recorded in the stratum radiatum of hippocampal slices from WT or 5xFAD mice perfused with 200 μM of indicated monosaccharide in aCSF. Scatter plots of fEPSP slope and representative traces before (black) and after (red) perfusion are shown. In contrast to L-fucose (*n* = 14), D-mannose (*n* = 4) (**a**) and D-galactose (*n* = 6) (**b**) did not enhance fEPSP. 5xFAD slices did not show enhanced fEPSP in response to L-fucose (*n* = 10) (**c**), D-galactose (*n* = 5) (**d**), or D-fucose (*n* = 5) (**e**).

**Supplementary Fig 3.**
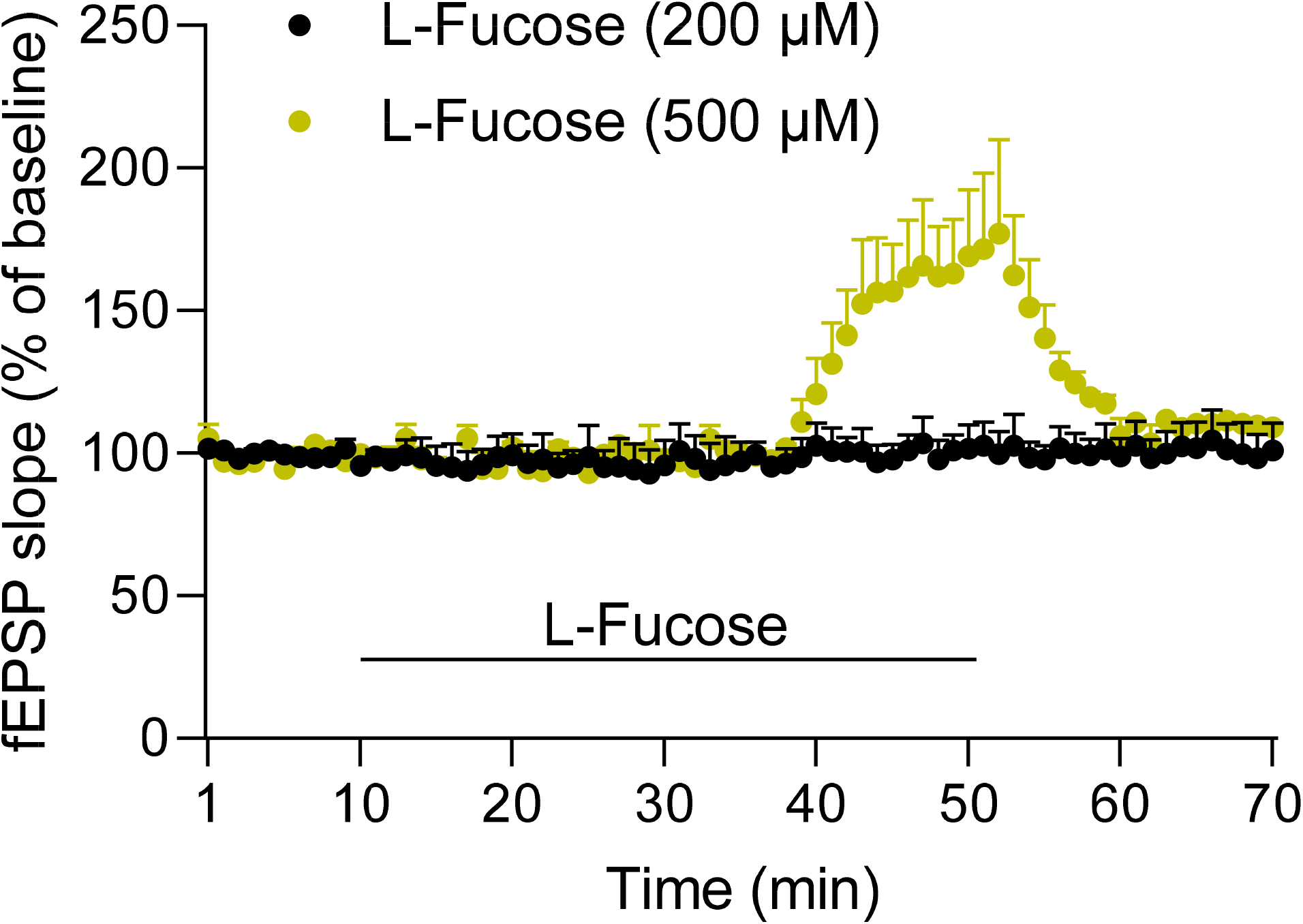
APP-PS1 hippocampus shows subdued responses to L-fucose. fEPSPs of Schaffer-collateral-CA1 synapses were recorded in the stratum radiatum of hippocampal slices from 12 months-old APP-PS1 mice perfused with 200 μM or 500 μM L-fucose in aCSF during indicated period. Shown are scatter plots of fEPSP slope; *N* = 4/group.

**Supplementary Fig. 4.**
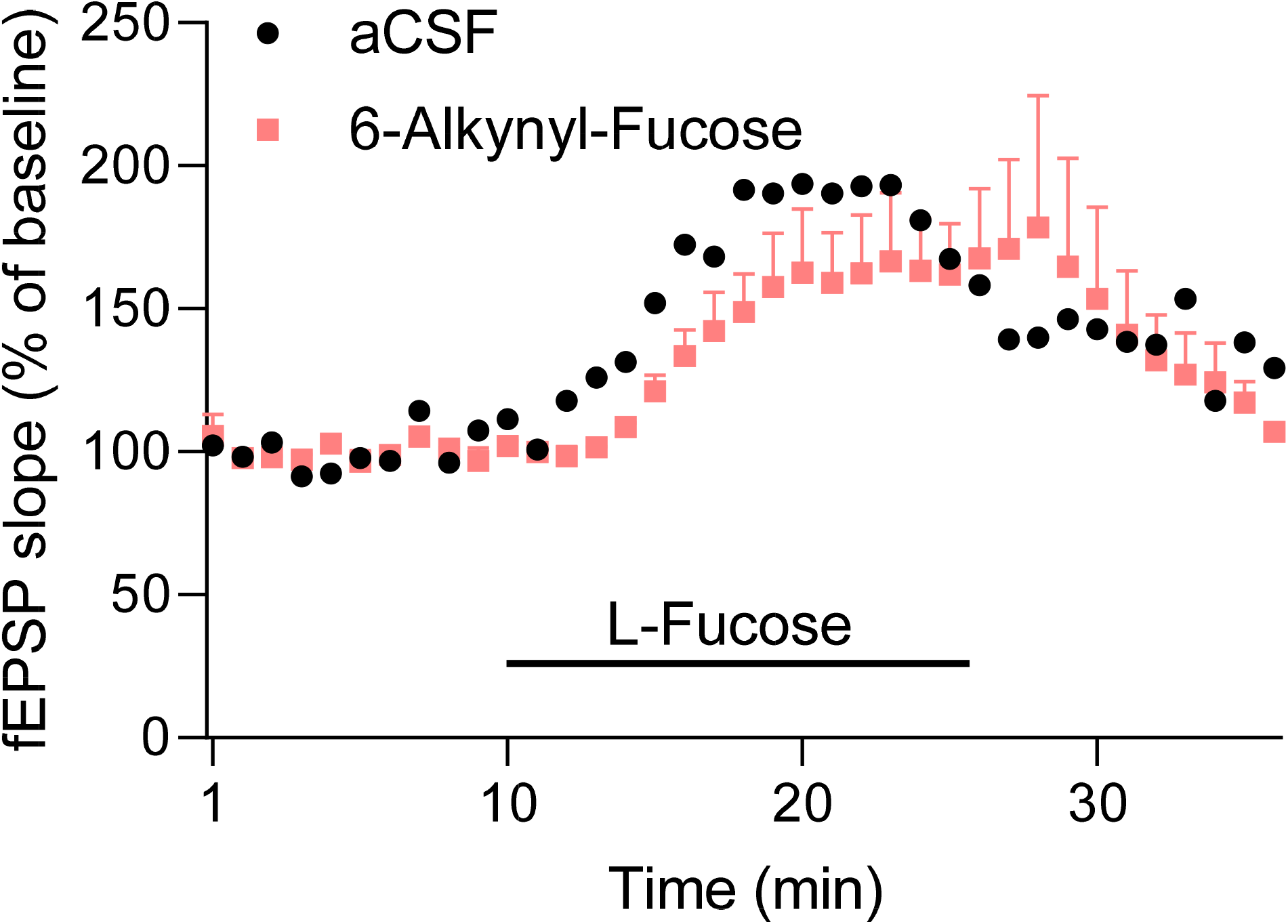
Inhibition of fucosylation does not affect responses to L-fucose. Scatter plots of fEPSP slope in WT slices with or without pre-incubation with 200 μM 6-alkynyl fucose, a fucosylation inhibitor, for 1 hr. *N* = 2/group.

**Supplementary Fig. 5.**
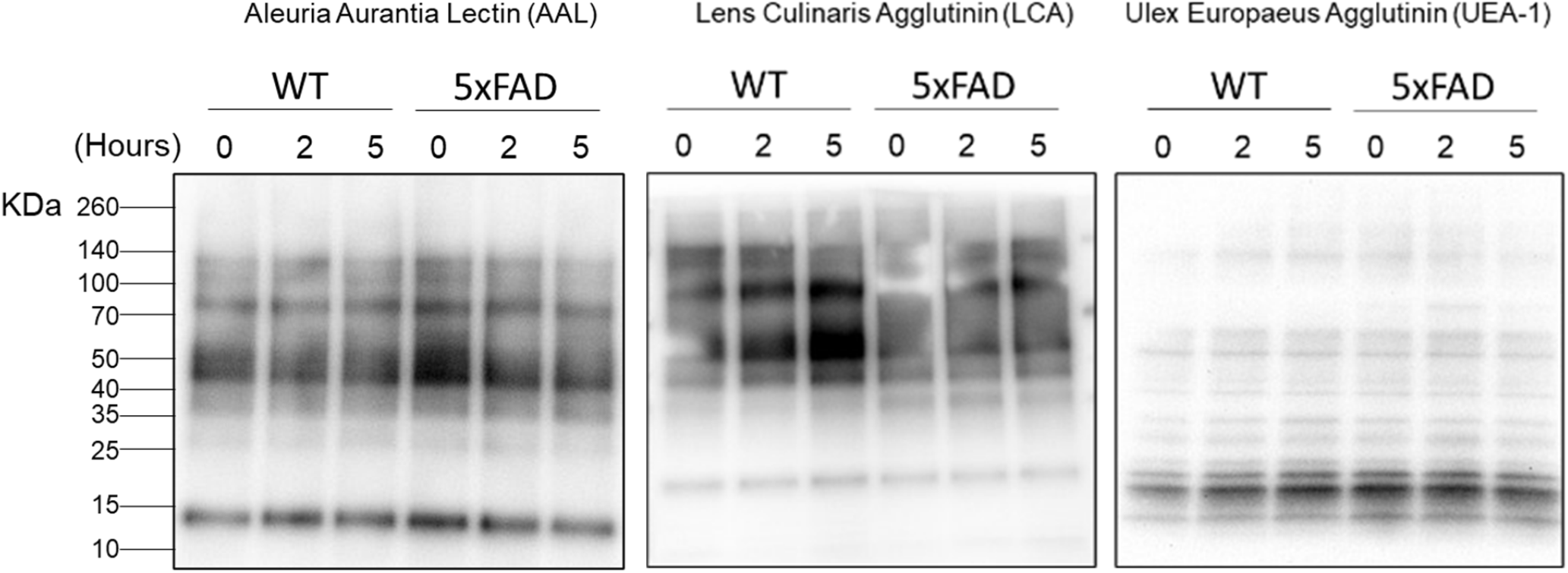
L-fucose treatment does not alter macromolecule fucosylation. Lectin blots of homogenates generated from WT or 5xFAD hippocampal slices perfused with 200 μM L-fucose for indicated hours. No significant alterations of reactivities were observed using the three indicated lectins recognizing fucosylated glycans.

**Supplementary Fig. 6.**
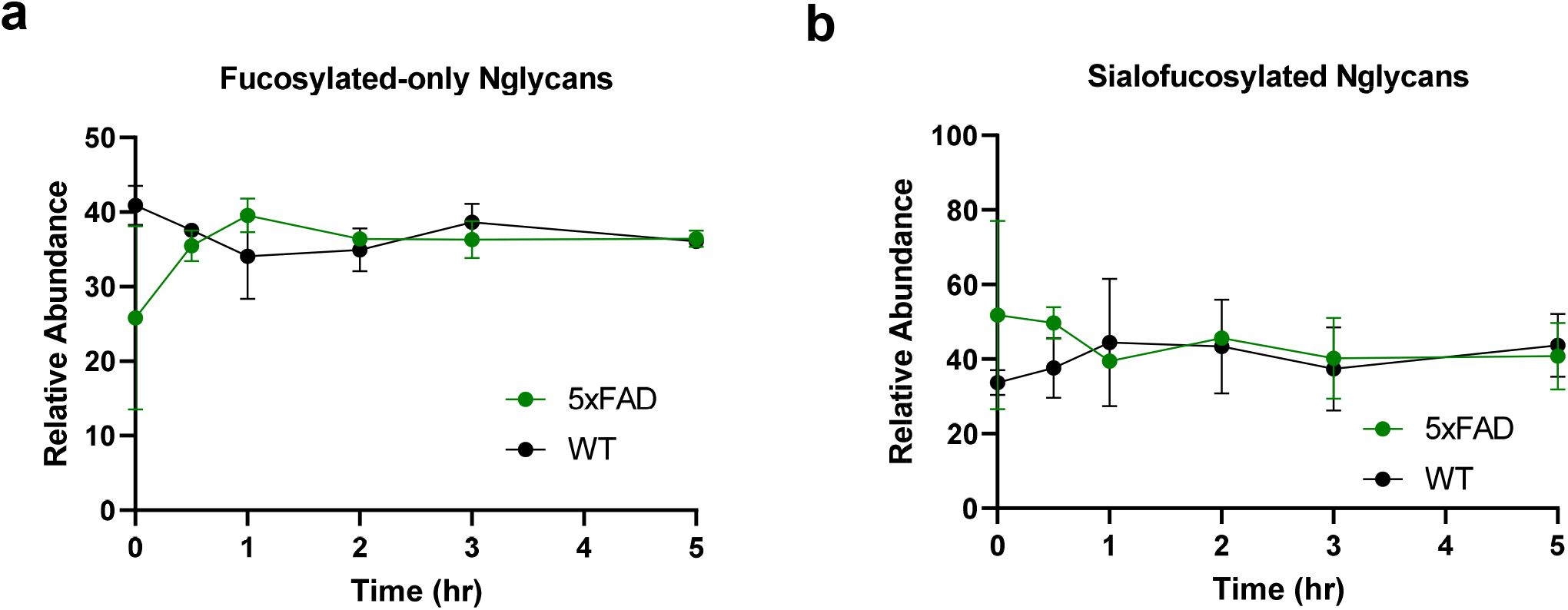
L-fucose treatment does not alter the relative abundance of fucosylated N-glycans. Shown are the relative abundances of fucosylated-only N-glycans (**a**) and sialofucosylated N-glycans (**b**) in WT hippocampal slices following perfusion of L-fucose (200 μM) for 0, 0.5, 1, 2, 3, and 5 hrs. *N* = 2 per time point/genotype. Data were extracted from comprehensive N-glycomics analyses (see online Methods). No statistically significant difference was observed between WT and 5xFAD mice nor between different time points.

**Supplementary Fig. 7.**
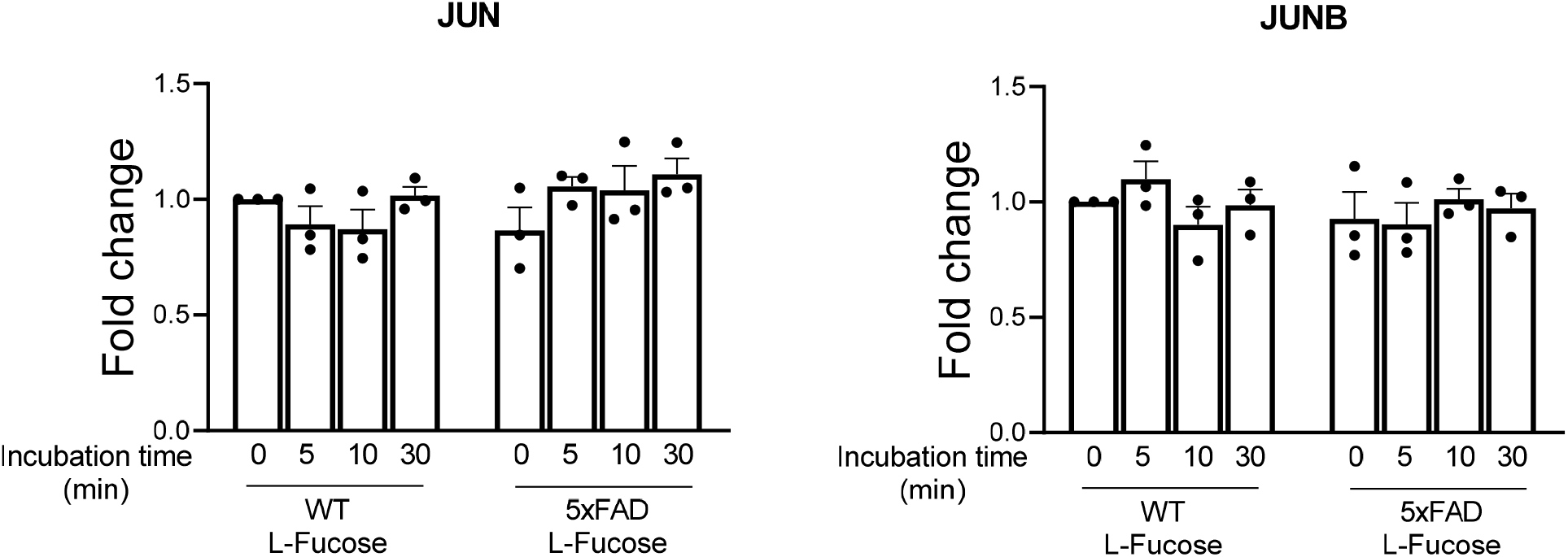
L-fucose effects on *c-fos* and *bdnf* is specific. In contrast to the time-dependent induction of *c-fos* and *bdnf* expressions by L-fucose shown in Fig. 4c, the same hippocampal samples did not show inductions of *jun* or *junB* expressions. *N* = 3/group.

**Supplementary Fig. 8.**
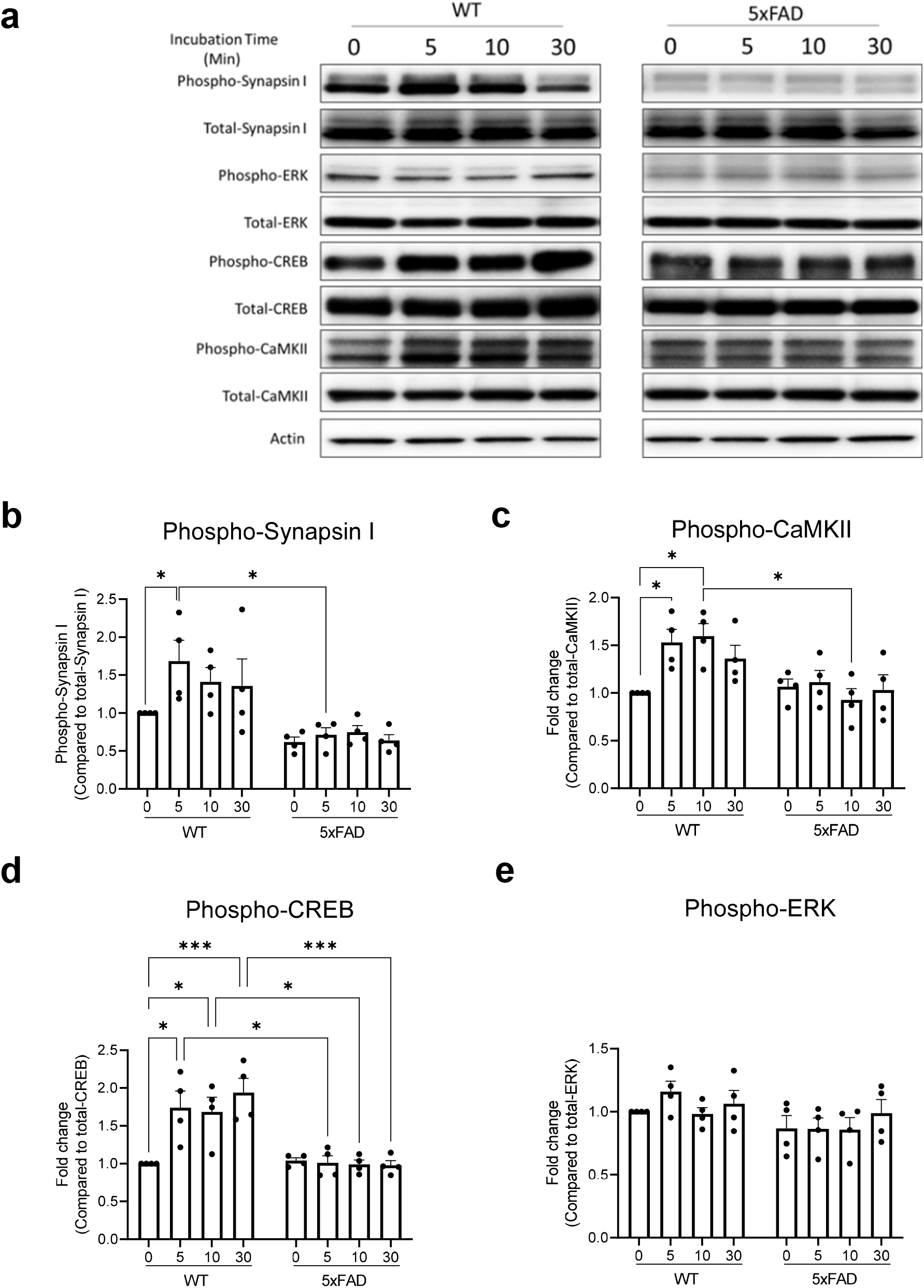
L-fucose rapidly activates presynaptic release-enabling signaling pathways. Western blotting of homogenates from hippocampal slices of WT and 5xFAD mice incubated with L-Fucose (200 μM) for 0, 5, 10, and 30 min. Shown are representative blots (**a**) and ratios of phosphorylated/total levels of indicated proteins based on band intensities (**b-e**), from four independent experiments. Two-way ANOVA with Tukey’s post hoc test. \**p*<0.05; \*\**p*<0.01; \*\*\**p*<0.001. The analysis shows significant differences between WT and 5xFAD in all four proteins - phospho-synapsin I: F(1)=28.48, *p*<0.0001; phospho-CaMKII: F(1)=39.20, *p*<0.0001; phospho-CREB: F(1)=15.48, *p*<0.001; and phospho-ERK: F(1)=6.59, *p*=0.017. The analysis also shows significant time-dependent differences in phospho-synapsin I: F(3)=3.64, *p*=0.048 and in phospho-CaMKII: F(3)=4.19, *p*=0.016.

**Supplementary Fig. 9.**
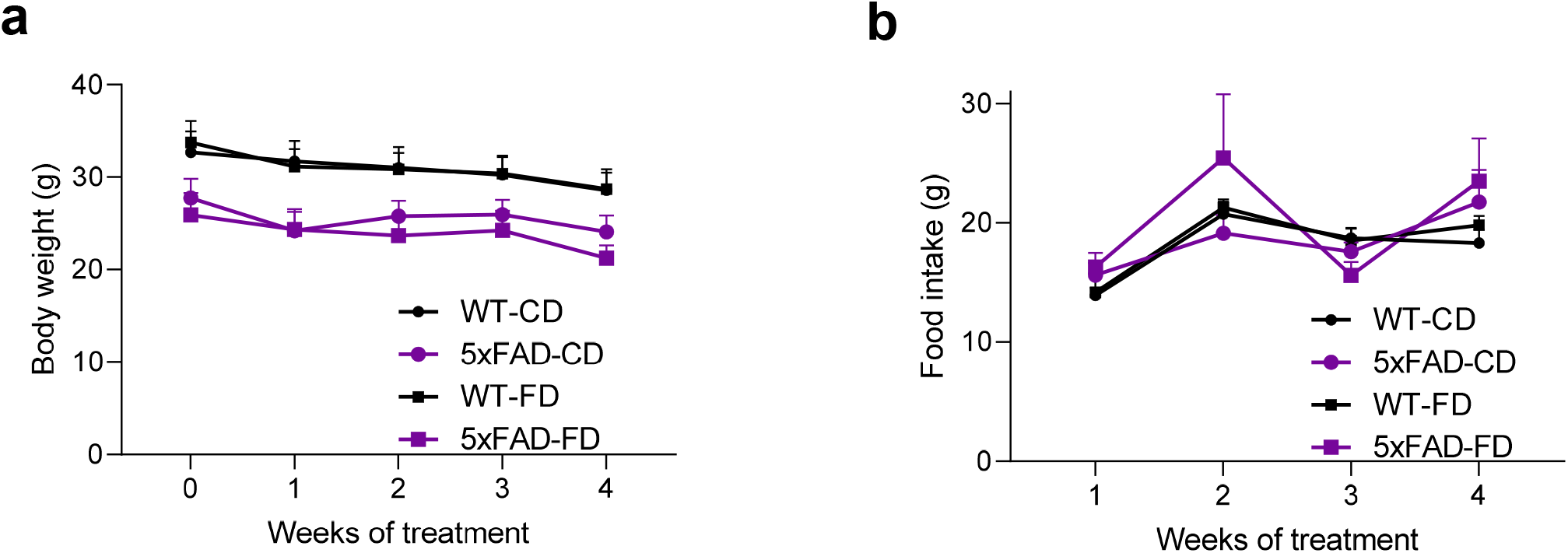
FD consumption does not affect body weight or food intake. Body weight (a) and food intake (b) of WT and 5xFAD mice during the month of receiving CD or FD. WT-CD, *n* = 11; 5xFAD-CD, *n* = 12; WT-FD, *n* = 13; and 5xFAD-FD, *n* = 11.

**Supplementary Fig. 10.**
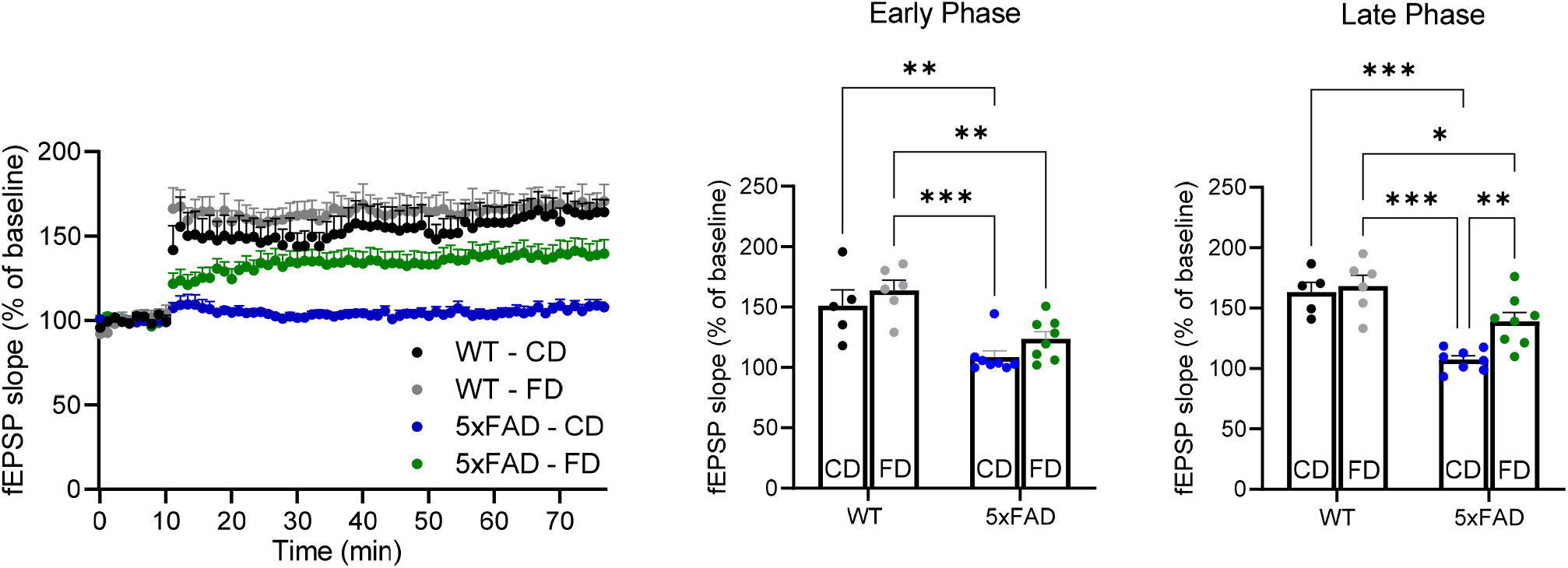
FD consumption improves hLTP in 5xFAD mice. Hippocampal LTP was recorded in WT and 5xFAD mice following one month treatment of CD or FD. Shown are representative tracings of hLTPand quantification of fEPSP slope during the early and late phases of hLTP. WT-CD, *n* = 6; 5xFAD-CD, *n* = 8; WT-FD, *n* = 6; and 5xFAD-FD, *n* = 8. \**p*<0.05, \*\**p*<0.01, \*\*\**p*<0.001; two-way ANOVA with Tukey’s post-hoc test.

**Supplementary Fig. 11.**
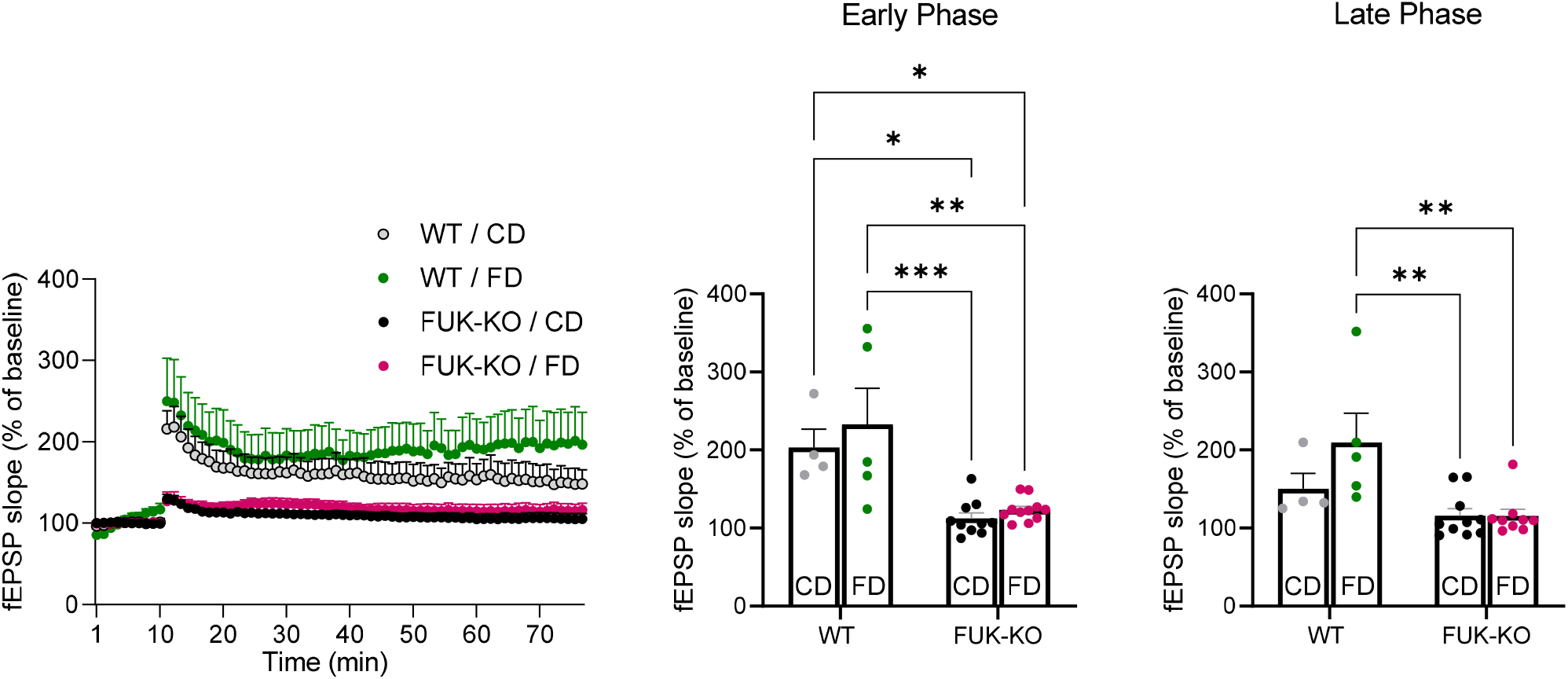
FD consumption does not improve hLTP deficits in FUK-KO mice. Hippocampal LTP was recorded in WT and FUK-KO mice following one month treatment of CD or FD. Shown are representative tracings of hLTPand quantification of fEPSP slope during the early and late phases of hLTP. WT/CD, *n* = 5; FUK-KO/CD, *n* = 10; WT/FD, *n* = 5; and FUK-KO/FD, *n* = 11. \**p*<0.05, \*\**p*<0.01, \*\*\**p*<0.001; two-way ANOVA with Tukey’s post-hoc test.

**Supplementary Fig. 12.**
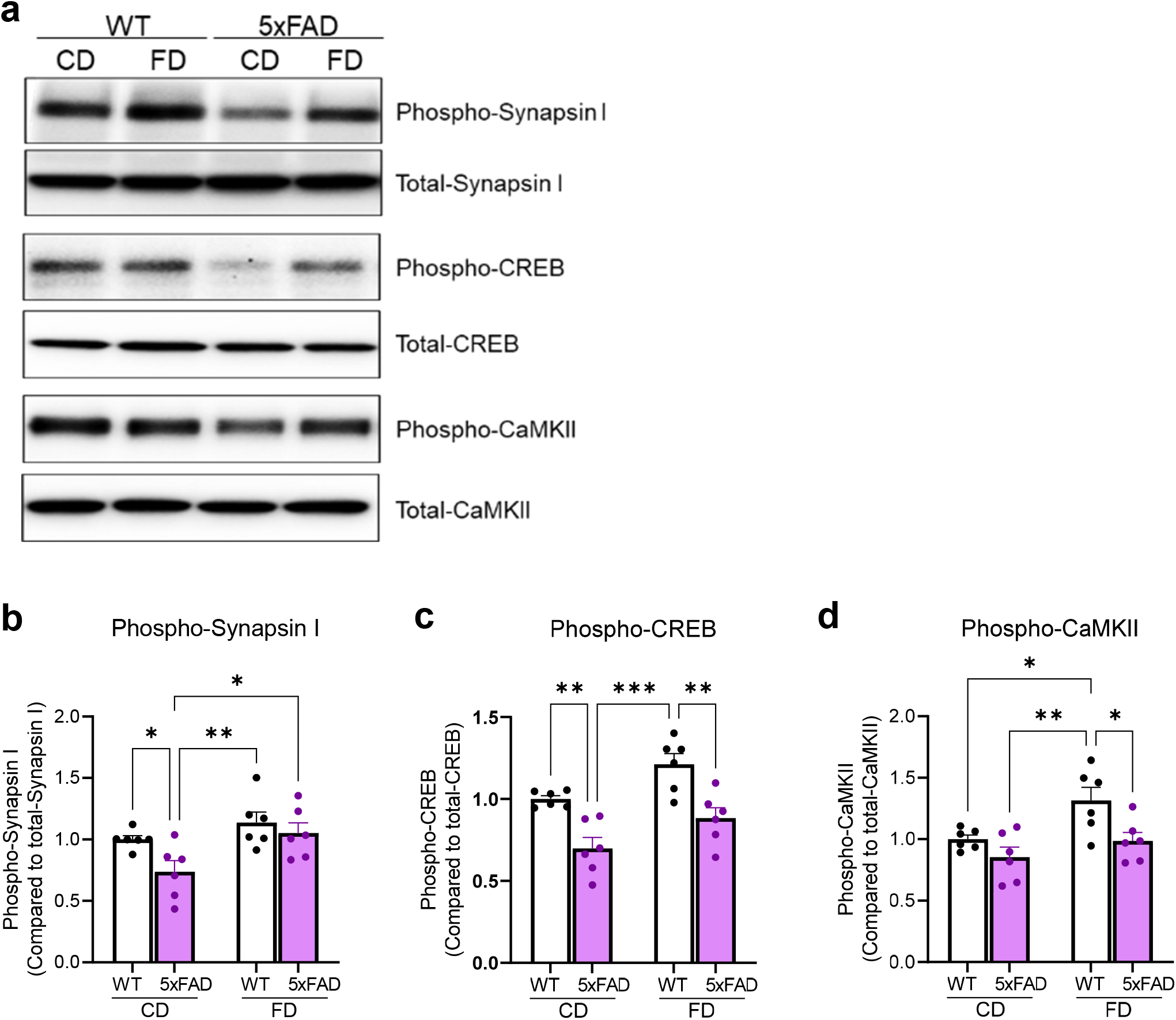
FD consumption activates molecules involved in synaptic plasticity. Western blotting of homogenates from hippocampi of WT and 5xFAD following one month consumption of CD or FD. Shown are representative blots (**a**) and ratios of phosphorylated/total levels of indicated proteins based on band intensities (**b-d**). *N*=6/group; two-way ANOVA with Tukey’s post hoc test. \**p*<0.05; \*\**p*<0.01; \*\*\**p*<0.001. The analysis shows significant differences between WT and 5xFAD in all three proteins – phospho-synapsin I: F(1) = 5.13, *p* = 0.035; phospho-CREB: F(1) = 29.81 p<0.00001; and phospho-CaMKII: F(1) = 9.42, *p* = 0.006. The analysis also shows significant differences between diet groups in all three proteins – phospho-synapsin I: F(1) = 8.73, *p* = 0.008; phospho-CREB: F(1) = 11.75, *p* = 0.003; and phospho-CaMKII: F(1) = 8.34, *p* = 0.009.

**Supplementary Fig. 13.**
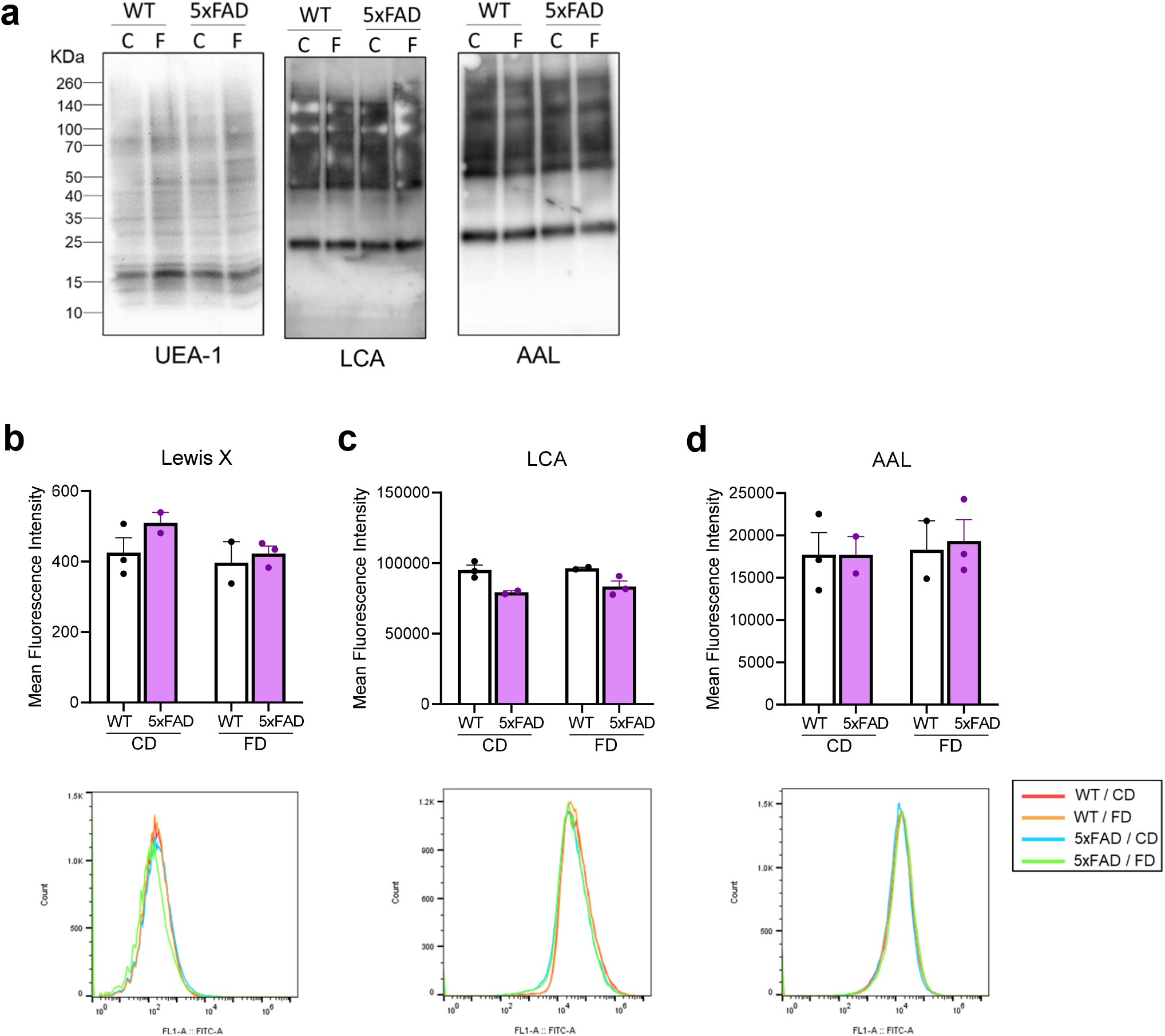
FD consumption does not alter macromolecule fucosylation in the brain. (**a**) Lectin blots of brain homogenates of WT or 5xFAD mice following one month of CD or FD consumption. No significant alterations of reactivities were observed using the three indicated lectins recognizing fucosylated glycans. (**b-d**) Brain cells were dissociated from fresh brain tissues, stained with the antibody to Lewis X antigen or the lectins LCA and AAL, and analyzed by flow cytometry. Shown are mean fluorescence intensities. WT-CD, *n* = 3; 5xFAD-CD, *n* = 2; WT-FD, *n* = 2; and 5xFAD-FD, *n* = 3.

**Supplementary Fig. 14.**
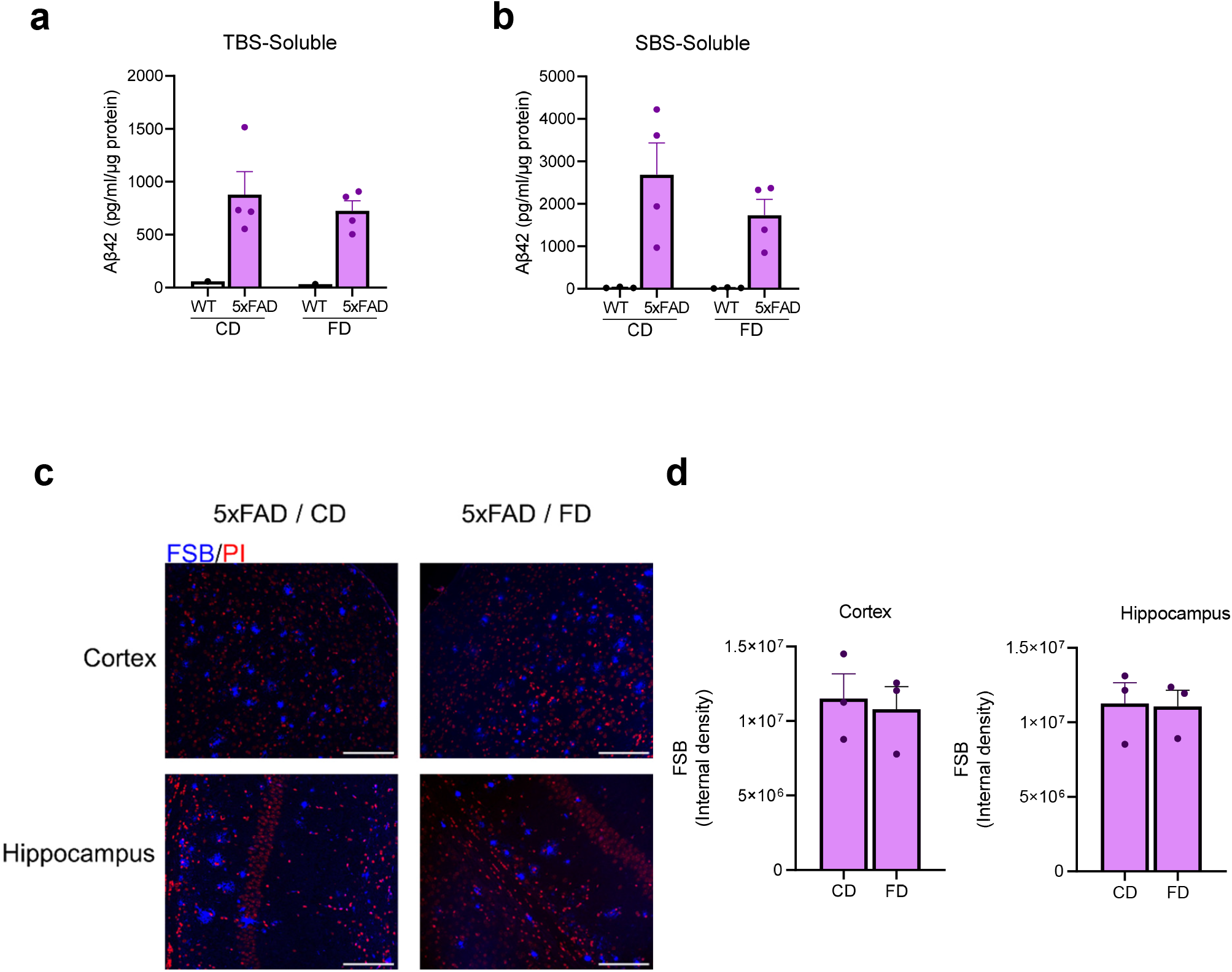
FD consumption does not affect Aβ amyloid pathology in 5xFAD mice. (**a-b**) Brain samples of WT or 5xFAD mice following one month of CD or FD consumption were differentially extracted into TBS-soluble (**a**) and SDS-soluble (**b**) fractions and their Aβ42 levels were quantified by ELISA. *N* = 4/group, Student’s unpaired t-test shows no statistical differences between groups. (**c-d**) Cortical and hippocampal sections were stained with the amyloid dye FSB and counterstained with PI. Shown are representative micrographs (**c**) (size bar = 100 μm) and quantification of FSB reactivities (**d**). *N* = 3/group, Student’s unpaired t-test shows no statistical differences between CD-fed and FD-fed 5xFAD mice groups.

**Supplementary Table 1.**
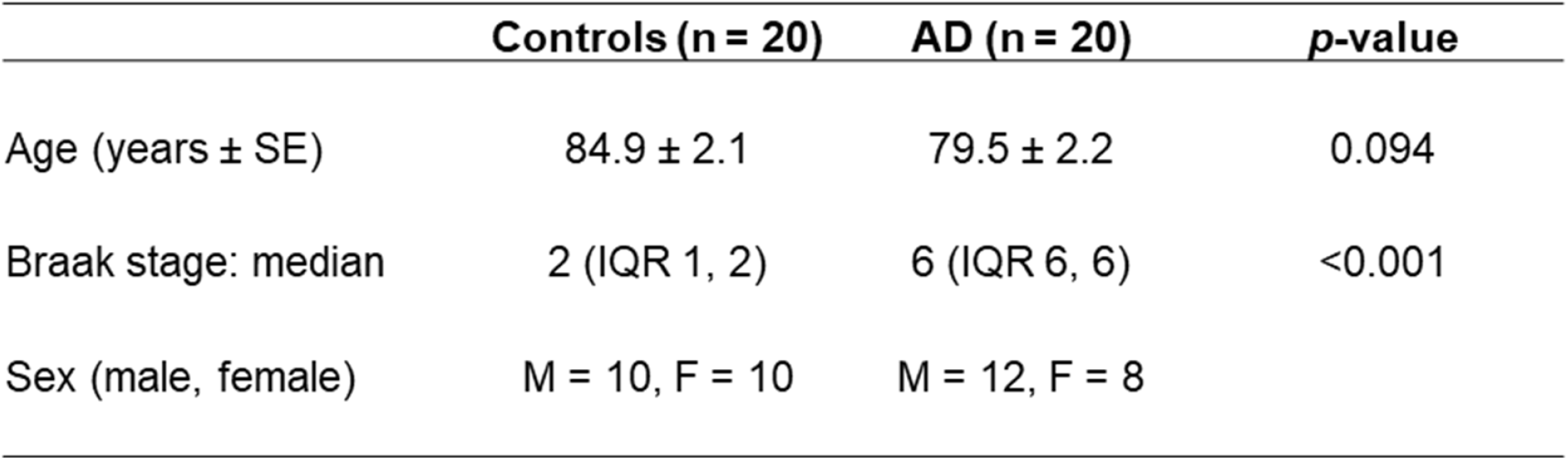

|  | **Controls (n = 20)** | **AD (n = 20)** | ***p*-value** |
| --- | --- | --- | --- |
| Age (years ± SE) | 84.9 ± 2.1 | 79.5 ± 2.2 | 0.094 |
| Braak stage: median | 2 (IQR 1, 2) | 6 (IQR 6, 6) | <0.001 |
| Sex (male, female) | M = 10, F = 10 | M = 12, F = 8 |  |

